# Mutations in Tomato ACC Synthase2 Uncover Its Role in Development beside Fruit Ripening

**DOI:** 10.1101/2020.05.12.090431

**Authors:** Kapil Sharma, Soni Gupta, Supriya Sarma, Meenakshi Rai, Yellamaraju Sreelakshmi, Rameshwar Sharma

## Abstract

The role of ethylene in plant development is mostly inferred from its exogenous application. The usage of the mutants affecting ethylene biosynthesis proffers a better alternative to decipher its role. In tomato, 1-aminocyclopropane carboxylic acid synthase2 (ACS2) is a key enzyme regulating ripening-specific ethylene biosynthesis. We characterized two contrasting *acs2* mutants; *acs2-1* overproduces ethylene, has higher ACS activity, and increased protein levels, while *acs2-2* is an ethylene under-producer, displays lower ACS activity, and protein levels than wild type. Consistent with high/low ethylene emission, the mutants show opposite phenotypes, physiological responses, and metabolomic profiles than the wild type. The *acs2-1* showed early seed germination, faster leaf senescence, and accelerated fruit ripening. Conversely, *acs2-2* had delayed seed germination, slower leaf senescence, and prolonged fruit ripening. The phytohormone profiles of mutants were mostly opposite in the leaves and fruits. The faster/slower senescence of *acs2-1*/*acs2-2* leaves correlated with the endogenous ethylene/zeatin ratio. The genetic analysis showed that the metabolite profiles of respective mutants co-segregated with the homozygous mutant progeny. Our results uncover that besides ripening, ACS2 participates in vegetative and reproductive development of tomato. The distinct influence of ethylene on phytohormone profiles indicates intertwining of ethylene action with other phytohormones in regulating plant development.

## INTRODUCTION

Ethylene is a simple gaseous molecule that also acts as a natural plant hormone. It participates in a multitude of development processes such as organ senescence, biotic and abiotic stresses, and fruit ripening (Abeles et al., 1992). The ethylene-mediated stimulation of the ripening process is restricted to climacteric fruits, where a surge in respiration marks the onset of ripening. The respiratory surge is preceded by increased ethylene biosynthesis that triggers ripening (Grierson, 2013). Antagonistically, mutations, or chemical treatments that block ethylene biosynthesis/perception, abolish or delay ripening of climacteric fruits (Brady, 1987; Martínez-Romero et al., 2007). For studies on climacteric fruit ripening, tomato (*Solanum lycopersicum*) has emerged as a preferred model system due to the availability of several monogenic mutants affecting the ripening process and ease of transgenic manipulations (Barry, 2014).

Studies carried out on tomato mutants defective in fruit ripening established a hierarchical genetic regulation of the ripening process. One of the extensively investigated mutants is *ripening-inhibitor* (*rin*) that lacks almost all ripening associated processes like the accumulation of lycopene, softening of fruits, and a climacteric burst of ethylene (Tigchelaar et al., 1978). The *RIN* gene encodes a MADs-type transcriptional factor, which positively regulates the onset of ripening process (Vrebalov et al., 2002). RIN directly binds to promoters of a large number of ripening-related genes, including those involved in ethylene biosynthesis (Fujisawa et al., 2012; Qin et al., 2012). The mutated *rin* gene encodes an in-frame fused RIN and Macrocalyx protein which strongly represses the expression of target genes (Ito et al., 2017). Consequently, the *rin* mutant does not ripen on exposure to exogenous ethylene, though it retains some ethylene-induced responses independent of RIN (Lincoln and Fischer, 1988).

The paramount role of ethylene in regulating tomato ripening is highlighted by the loss of ripening in tomato *Nr* mutant. The *Nr* mutant encodes a truncated ethylene receptor ETR3, resultantly compromised in ethylene perception (Wilkinson et al., 1995). The ripening can be restored in transgenic *Nr* fruits by antisense inhibition of mutated *ETR3* gene suggesting receptor inhibition model of ethylene action (Hackett et al., 2000). Likewise, overexpression of *EIL1* (*EIN3*-like transcription factor) restores ripening in *Nr* (Chen et al., 2004), indicating the operation of normal ethylene signal transduction in the mutant. The mutations in *ETR1*, *ETR4*, and *ETR5* genes had a differential effect on tomato ripening, probably by affecting their ethylene sensitivity (Okabe et al., 2011; Mubarok et al., 2019).

The elegant studies initially conducted on wounded apple fruit tissues uncovered the ethylene biosynthesis pathway in higher plants named as Yang cycle (Adams and Yang, 1979). The ethylene is derived from C-3,4 of amino acid methionine, which is first converted to S-adenosyl-L-methionine (SAM) by SAM synthase. SAM, a common precursor of many biosynthetic pathways, is converted to 1-aminocyclopropane carboxylic acid (ACC) by ACC synthase (ACS). The conversion of ACC to ethylene is catalyzed by ACC oxidase (ACO) in the presence of oxygen. The formation of ACC by ACS constitutes the first committed step in ethylene biosynthesis.

ACS is considered as the rate-limiting enzyme in the ethylene biosynthesis pathway. In tomato, ACS is encoded by a multigene family consisting of at least nine genes (*ACS1A, ACS1B*, and *ACS2-8*) and five putative genes (Liu et al., 2015). During tomato fruit development and ripening, *ACS* genes are differentially regulated by endogenous mechanisms and ethylene-mediated autocatalytic induction. During tomato fruit development, expressions of *ACS* genes considerably differ in system-I and system-II responses (Klee and Giovannoni, 2011). During system-I that is confined to the fruit expansion phase, *ACS* genes are moderately expressed. During system-II, which marks ripening induction, *ACS2, ACS4*, and *ACS6* are highly expressed (Barry et al., 1996, 2000; Van de Poel et al., 2012). The strong association between ripening and increased *ACS2* expression was indicated by the total suppression of tomato fruit ripening by antisense inhibition of the *ACS2* gene (Oeller et al., 1991). The high expression of *ACS2* during ripening likely results from the binding of transcription factors like RIN (Ito et al., 2008) and TAGL1 (Itkin et al., 2009) to the promoter of *ACS2* gene. Genome-wide analysis of DNA methylation demonstrated that tomato ripening is associated with the demethylation of several promoters, including *ACS2* and ethylene receptors *ETR3* and *ETR4* (Zhong et al., 2013). Based on the absence of binding of TAGL to *ACS4* promoter, Itkin et al. (2009) suggested *ACS2* is the primary ethylene biosynthesis gene, regulating tomato ripening. The kinetic analysis of ethylene biosynthesis, ACC accumulation, ACS activity, and *ACS* genes expression indicated that during tomato ripening ACS4 and ACS6 are less important, whereas ACS2 seems to be the main biosynthetic enzyme as its expression is ethylene dependent and mainly occurs during system 2 (Van de Poel et al., 2014).

While the *ACS2* gene along with other ethylene biosynthesis genes, seemingly contributes to the climacteric rise of ethylene during tomato ripening, information about its role in other developmental processes of tomato is limited. The paucity of information about its role stems from the near absence of tomato mutants compromised in the ethylene biosynthesis pathway. Here we describe isolation and characterization of two novel *ACS2* mutants of tomato having diametrically opposite effects on ethylene emission. The *acs2-1* mutant has high ethylene emission, and *acs2-2* shows reduced ethylene emission. We show that stimulation/reduction of ethylene emission affects several developmental processes right from seed germination to fruit ripening. We also show that variation in ethylene emission affects hormonal and metabolome profiles in an opposite manner.

## RESULTS

### Isolation of *acs2* mutants

To identify *ACS2* gene mutants, we screened genomic DNA from 9,144 EMS-mutagenized M_2_ plants by TILLING. In total, nine alleles were identified, of which two named *acs2-1* (M82-M_3_-112) and *acs2-2* (M82-M_2_-162A) were characterized in detail **(Supplemental Table S1)**. Sequencing of the full-length gene, including promoter, revealed two exonic and three intronic mutations in *acs2-1*, and one intronic and two promoter-localized mutations in *acs2-2* **(Figure 1A, Supplemental Table S2)**. Homozygous mutant lines were characterized in M_6_ generation and backcrossed twice to parent M82 plants. Genetic segregation analysis showed that *acs2-1* and *acs2-2* were inherited as a monogenic Mendelian trait. Segregation of mutation in F_2_ progeny was monitored by CEL-I endonuclease assay (Mohan et al., 2016) [*acs2-1* BC_1_F_2_-120-plants, 29 (*acs2-1*/*acs2-1*), 61 (*ACS2*/*acs2-1*), and 30 (*ACS2*/*ACS2*), ratio 1:2:1, χ^2^ (0.224) P = 0.62]; [*acs2-2* BC_1_F_2_-total 108-plants, 25 (*acs2-2*/*acs2-2*), 56 (*ACS2*/*acs2-2*), and 27 (*ACS2*/*ACS2*), ratio 1:2:1, χ^2^ (0.101) P = 0.95] **(Supplemental Table S3)**. For the majority of the fruit ripening experiments, we compared respective mutants with parental wild type (WT), and BC_1_F_2_ progeny.

**Figure 1.**
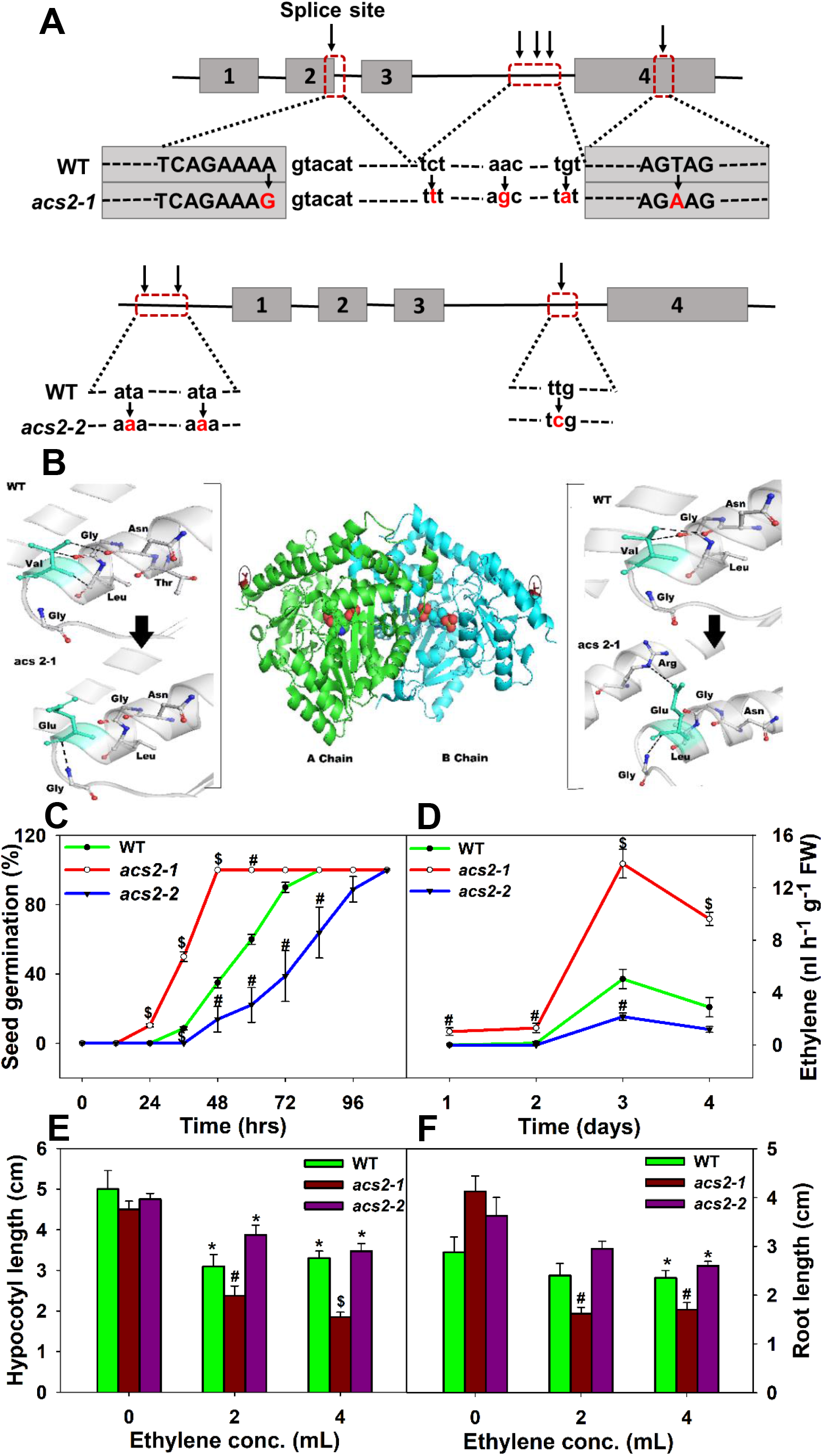
Characterization of *acs2* mutants. **(A)** Localization of mutations in the *ACS2* gene in *acs2–1* and *acs2–2*. Arrows indicate the base changes in the introns, exons, splice site, and promoter. (**B**) Model of ACS2 protein comprising of A and B chains. The altered intermolecular bonding in the respective chain of ACS2 protein due to V352E mutation in *acs2–1* is depicted in the side panels (Red arrows). **(C)** Time course of mutant and WT seed germination (n = 30 ± SE). **(D)** Time course of ethylene emission from dark-grown *acs2* mutants and WT seedlings. At different time points from sowing, seedlings were transferred to an airtight vial for 24 h for estimation of ethylene. **(E, F)** Effect of exogenous ethylene on the root (**E**) and hypocotyl elongation (**F**) of 5-day-old dark-grown seedlings. Unless otherwise mentioned, the statistical significance marked in all figures was determined using Student’s *t*-test. (* for P ≤ 0.05, # for P ≤ 0.01 and $ for P ≤ 0.001).

### *In silico* analysis predicted over-expression/reduced-expression for *acs2* mutants

Splice site analysis by NetGene2-server (http://www.cbs.dtu.dk/services/NetGene2/) predicted that A398G (K100=) mutation located in *acs2-1* at 5′ splice site terminating the second exon leads to more efficient mRNA splicing that in turn may enhance *acs2-1* transcript level **(Supplemental Figure S1)**. Computational protein modeling and protein stability analysis indicated that V352E change affects bonding and folding pattern, improving the stability of ACS2-1 protein (**Figure 1B**, **Supplemental Table S4–5).** Taken together, enhanced transcript level and stable ACS2-1 protein may lead to an over-expression hypermorphic mutation.

It is reported that C(A/T)_8_G sites also have an affinity for RIN binding (Fujisawa et al., 2013). In *acs2-2*, T-106A mutation is located in C(A/T)_9_G location, therefore it is unlikely that it may have disrupted RIN binding site. However, T-106A mutation disrupts a SOC1 binding site and gains an AZF binding site (Kodaira et al., 2011; tomato homolog Solyc04g077980.1), a C2H2-type transcription factor negatively regulating ABA-mediated responses (**Supplemental Table S6, Supplemental dataset S1**). T-382A mutation is located at a methylated CpG site that shows progressive methylation reduction during ripening (http://ted.bti.cornell.edu/cgi-bin/epigenome/home.cgi) (**Supplemental Table S7, Supplemental dataset S2**). It is plausible that the above mutations reduce the affinity of regulatory transcription factors, thus compromising *acs2-2* gene expression resulting in a reduced-expression phenotype.

### *acs2-1* shows faster seed germination

In *acs2-1*, the onset of seed germination was 12h earlier than WT, whereas it was delayed by 12h in *acs2-2.* The attainment of full germination followed the above pattern, with *acs2-1* attaining 50% germination at 35h, WT at 55h, and *acs2-2* at 77h **(Figure 1C)**. Ethylene emission from respective mutant seedlings also showed a similar pattern. Three-day old *acs2-1* seedlings showed higher emission (280%), and *acs2-2* lower emission (40%) than WT **(Figure 1D)**. Mutant seedlings showed a similar pattern in ethylene-mediated growth inhibition with *acs2-1* being more sensitive and *acs2-2* less sensitive than WT **(Figure 1E-F)**.

### *acs2* mutants have contrasting phenotypes

During vegetative growth, *acs2-1* displayed faster growth, elongated internodes, and higher ethylene emission from detached leaves than WT (**Figure 2A,C**). In contrast, *acs2-2* exhibited slightly slower growth and lower ethylene emission than WT (**Figure 2B,C**). Akin to ethylene emission, detached mutant leaves showed faster (*acs2-1*) and slower (*acs2-2*) loss of coloration than WT **(Figure 2D)**. In mutant leaves, chlorophylls and carotenoids levels similarly differed from WT **(Supplemental Figure S2)**. Hormonal profiling of *acs2-1* leaves showed up-regulation of jasmonic acid (JA), ABA, methyl jasmonate (MeJA), and down-regulation of IAA, IBA, salicylic acid (SA), and zeatin than WT. In contrast, the hormonal profile of *acs2-2* was closer to WT, barring lower levels of SA, and higher levels of zeatin **(Figure 2E,F)**. PCA of primary metabolites of mutants was distinctly different from WT **(Figure 2G, Supplemental dataset S3)**. In general, more metabolites were downregulated in *acs2-1* than *acs2-2* **(Figure 2H)**. Key TCA cycle metabolites, such as citrate, isocitrate, and malate, showed opposite levels indicating a differential effect of *acs2-1* and *acs2-2* on the TCA cycle. In *acs2-2* levels of several amino acids were high, whereas, in *acs2-1* barring threonine, other amino acids showed lower abundance than WT.

**Figure 2.**
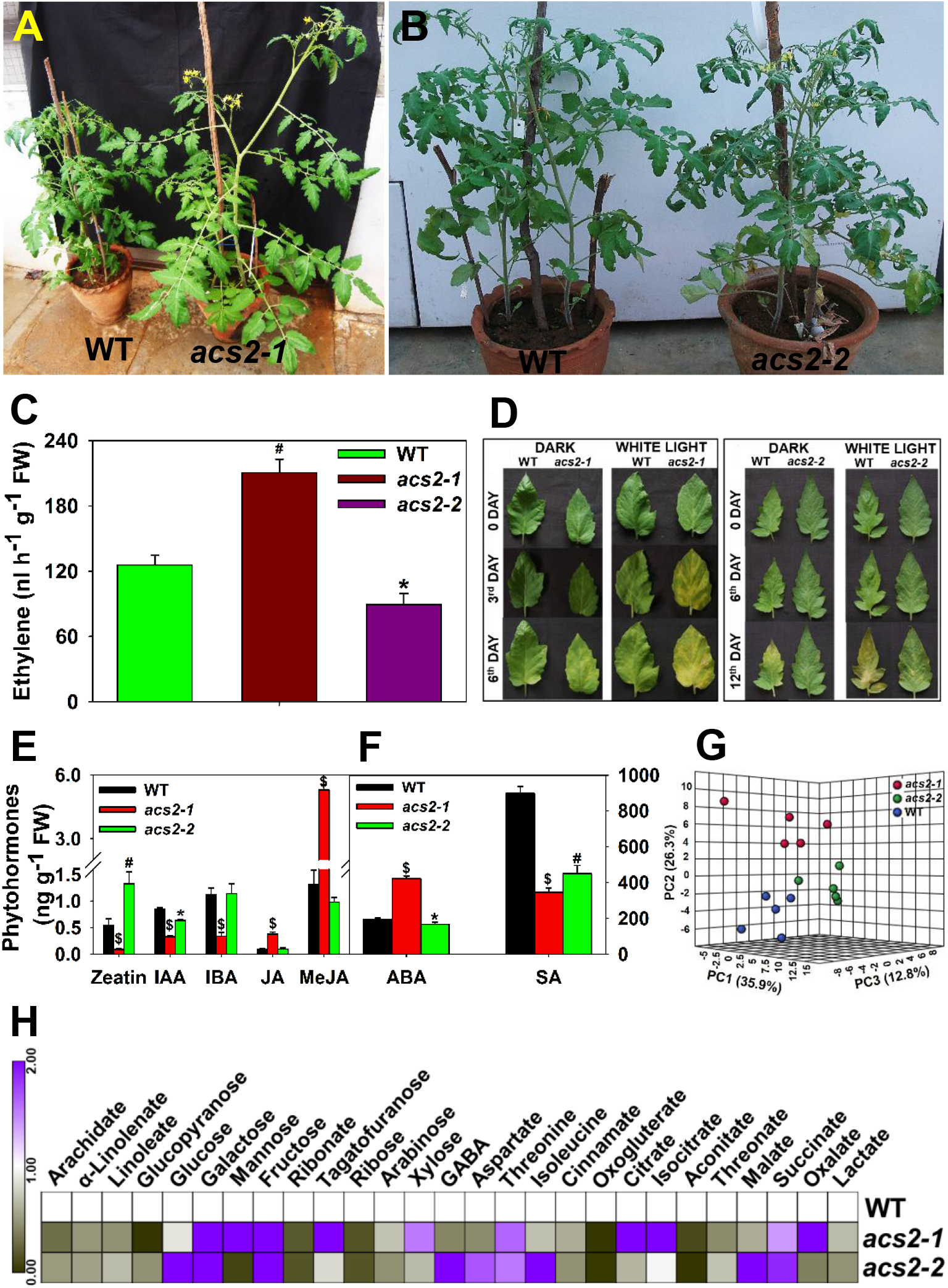
Characterization of *acs2* mutants and WT plants. **(A, B)** Phenotypes of greenhouse-grown WT and *acs2–1* (**A**) and *acs2–2* (**B**) plants. (**C**) Ethylene emission from leaves of WT and *acs2* mutants. (**D**) Senescence of detached leaves of WT and *acs2* mutants incubated in darkness or under the light. **(E, F)** Phytohormones profiles of WT and *acs2* mutants leaves. (**G**) Principal component analysis (PCA) of WT and *acs2* mutants (n = 5). The PCA was plotted using the MetaboAnalyst 4.0. (**H**) Heat maps showing significantly up- or down-regulated (> 1.5 fold) metabolites in *acs2* mutants with reference to WT. The relative changes between mutant and WT were calculated by determining the mutant/WT ratio for individual metabolites. The leaves harvested from the seventh node of 45-day-old plants were used for all analyses.

### *acs2-1* shows accelerated fruit ripening

Both *acs2* mutants also affected the reproductive phase. *acs2-1* had more flowers with a higher fruit set per truss, whereas *acs2-2* had fewer flowers and a lower fruit set per truss than WT (**Figure 3A**, **Supplemental Table S8**). These results indicate that *acs2-1* accelerates the overall reproductive phase and enhances fruit yield, and *acs2-2* has the opposite phenotype. Ripened fruits of *acs2-1* were smaller, and *acs2-2* were bigger than WT (**Figure 3B).** *acs2-1* fruits showed accelerated development reaching *m*ature *g*reen (MG) stage by 31 *d*ays *p*ost*-a*nthesis (dpa) **(Figure 3B,C)**. Post-MG stage, *acs2-1* showed accelerated ripening attaining *r*ed *r*ipe (RR) stage by 38 dpa. RR stage of *acs2-1* fruits was severely short (8-days) with the onset of fruit senescence (FS) at 46 dpa. Though WT and *acs2-2* fruits reached the MG stage at the same time, the duration to attain the RR stage was significantly longer in *acs2-2* **(Figure 3C,D)**. Post-RR stage, the onset of FS was also significantly delayed in *acs2-2* (40-days) than WT (25-days). While ripening *acs2-1* fruits emitted more ethylene, *acs2-2* fruits had lower ethylene emission than WT. Respiratory CO_2_ emission from *acs2-1*, *acs2-2*, and WT fruits followed a pattern similar to ethylene, indicating a linkage between these two processes **(Figure 3E-F)**. Analysis of homozygous backcrossed progeny of *acs2-1* and *acs2-2* showed ethylene emission and ripening pattern, nearly similar to the respective mutant (**Figure 3 C-E)**. *acs2-1* fruits were less firm at TUR and RR, while *acs2-2* fruits were more firm than WT. *acs2* mutations also had a minor effect on °Brix and pH value of fruits **(Supplemental Figure S3)**. The analysis of BC_4_F_2_ progeny of *acs2-1* and *acs2-2* in Arka Vikas (AV) and Pusa Early Dwarf (PED) cultivars also showed ethylene emission and ripening pattern, similar to respective mutant **(Supplemental Figure S4)**.

**Figure 3.**
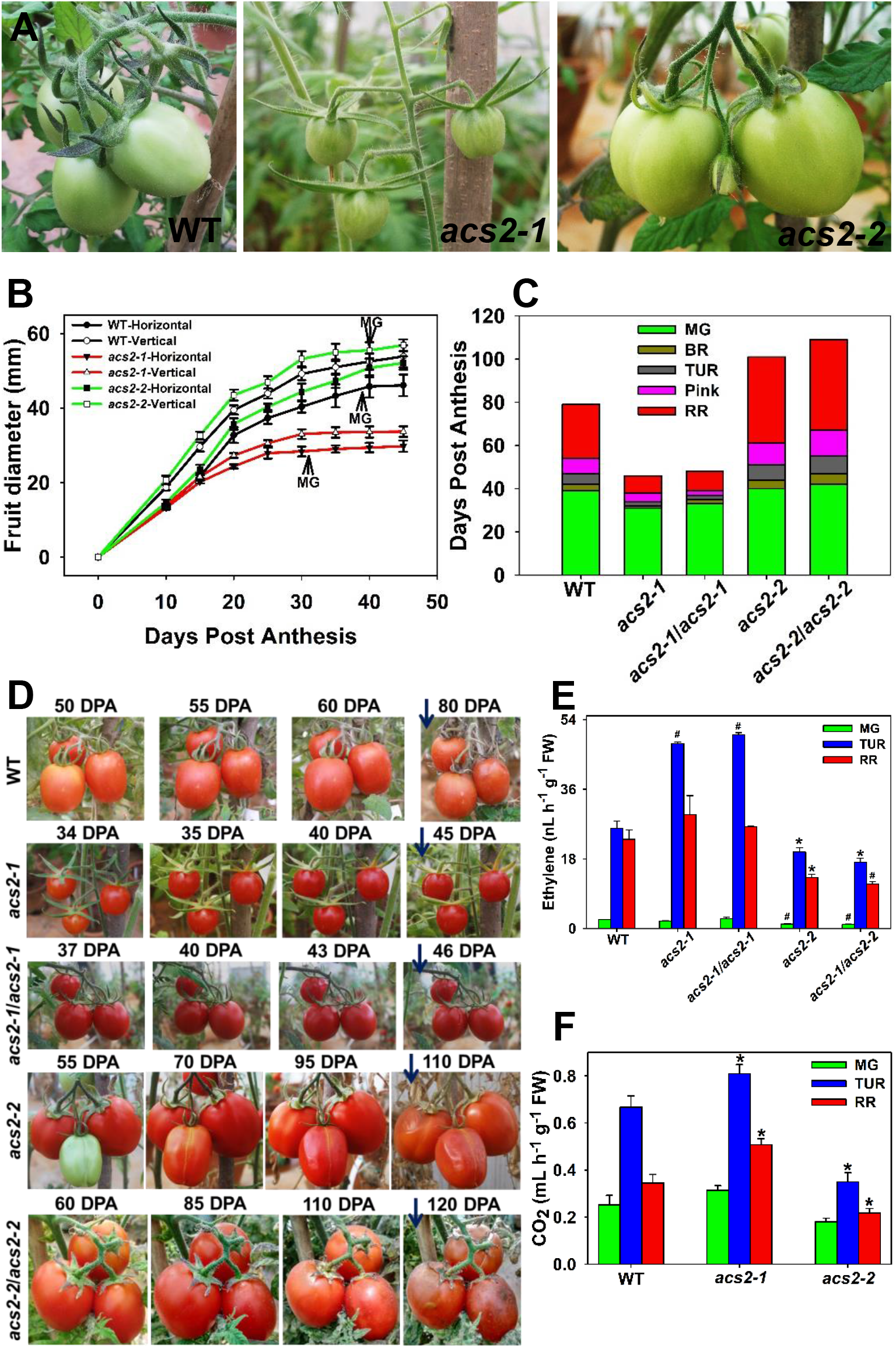
The development and on-vine ripening of *acs2* mutants and WT fruits. (**A**) WT and *acs2* mutant fruits (WT: 39 DPA, *acs2–1*: 30 DPA, *acs2–2*: 40 DPA). (**B**) Time course of the increase in fruit diameter from anthesis until 45 days. Arrows mark the attainment of the MG stage (n = 3). (**C**) Chronological development of tomato fruits post-anthesis until the onset of senescence. The stacked bar graph shows the attainment of different stages of ripening: mature green (MG), breaker (BR), Turning (TUR), Pink (P), and red ripe (RR) stages until senescence onset. The time point to attain specific ripening phases was visually monitored. The appearance of wrinkles on fruit skin was taken as time point of senescence onset (WT-82 DPA; *acs2–1* - 42 DPA; *acs2–1*/*acs2–1* - 44 DPA; acs2*-2* - 102 DPA; *acs2–2*/*acs2–2*-112 DPA) (**D**) On-vine ripening of second truss WT and *acs2* mutant fruits from RR until senescence onset (Arrowheads). (**E**) Ethylene emission from MG, TUR, and RR fruits of WT, *acs2* mutants, and their respective backcrossed homozygous progeny (*acs2–1*/*acs2–1; acs2–2*/*acs2–2*). (**F**) The CO_2_ emission from MG, TUR, and RR fruits of WT and mutants.

### *acs2* mutants differently affect ACS activity

In plants, ethylene synthesis is a two-step process, wherein SAM is converted to ACC by ACS. In the next step, ACO converts ACC to final end-product ethylene (Yang and Hoffman, 1984). We determined whether higher/lower ethylene emission from *acs2* fruits correlated with *in vivo* levels of ACC. Consistent with higher ethylene emission in *acs2-1*, the ACC level was higher, whereas, in *acs2-2*, it was lower than WT **(Figure 4A)**. *In vivo*, ACC is also conjugated to malonic acid to form malonyl-ACC (MACC) by ACC malonyltransferase (Hoffman et al., 1982). Akin to higher ACC, MACC level was also higher in *acs2-1*, whereas *acs2-2* had lower MACC levels than WT **(Figure 4B)**. To ascertain whether high/low ACC levels in mutants reflected in ACS enzyme activity, we *in vitro* assayed ACS activity. Congruent to ACC levels, ACS activity was 2-fold higher in *acs2-1*, and 0.5-fold in *acs2-2* compared to WT **(Figure 4C)**. Next, we examined whether high/low ACC levels also influenced the activity of the subsequent enzyme-ACO. Even ACO activity was significantly higher in *acs2-1*, whereas *acs2-2* had lower ACO activity **(Figure 4D)**.

**Figure 4.**
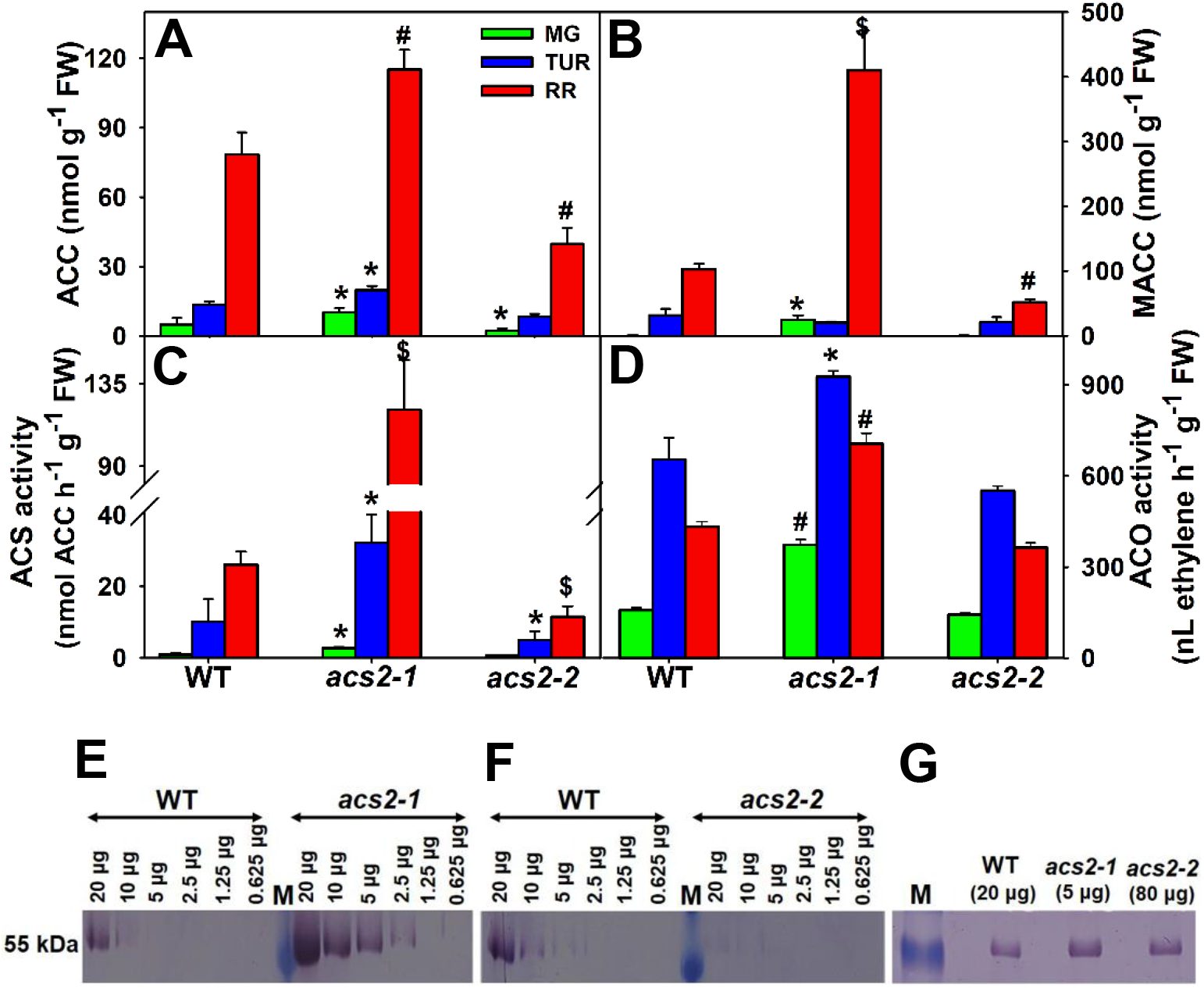
ACS activity and protein levels in mutants and WT fruits. **(A, B)** ACC (**A**) and MACC (**B**) levels in WT and *acs2* mutants. (**C, D**) ACS (**C**) and ACO (**D**) enzyme activities in WT and *acs2* mutants. (**E, F)** Comparison of ACS2 protein levels in RR fruits of WT, *acs2–1* (**E**), and *acs2–2* (**F**) using Western blotting. A similar amount of immunoprecipitated proteins from WT and *ACS2* mutants was serially diluted. Dilutions demonstrate that the abundance of ACS2 protein in *acs2–1* is 4-fold higher, and in *acs2–2*, it is 4-fold lower compared to WT. (**G**) The equal amount of ACS2 protein (Calculated based on data in **E** and **F**) loaded on Western blot shows similar staining with the Phospho-Ser antibody. **M** represents the protein marker

### *acs2* mutants differ in ACS protein levels

To determine whether altered ACS activity reflected changes in ACS2 protein levels, we raised polyclonal antibodies against ACS2 using a synthetic peptide, and antibody specificity was confirmed against the peptide **(Supplemental Figure S5)**. In ripening tomato pericarp, the level of ACS protein is estimated to be <0.0001% of total soluble protein (Bleecker et al., 1986). Due to its low abundance, we could not detect ACS2 protein on Western blot on the direct loading of supernatants from fruit homogenates. Therefore, we purified IgG fraction and immunoprecipitated ACS2 protein to enrich it before Western blot analysis **(Supplemental Figure S6).** Reduction in ACS activity in the supernatant after immunoprecipitation indicated the specificity of antibodies toward ACS2 protein. Western blot analysis revealed no discernible ACS2 protein band in MG and TUR fruits of *acs2-1, acs2-2*, and WT. Only in RR fruits, a single band of 55 kD was detected corresponding to the reported molecular size of tomato ACS2 protein in *acs2-1* and WT (Rottmann et al., 1991).

A more prominent band in *acs2-1* compared to WT indicated that it is enriched in ACS2 protein. In contrast, a very faint band in *acs2-2* RR fruits indicated a highly reduced ACS2 protein level **(Supplemental Figure S6)**. To ascertain the relative amount of ACS2 protein in acs2*-1, acs2-2*, and WT RR fruits, we subjected samples to serial dilution. Matching the intensity of bands of serially diluted proteins indicated that the level of ACS2 protein in WT was about 4-fold less than *acs2-1*, whereas ACS2 protein level in *acs2-2* was about 4-fold lower than WT **(Figure 4E-F)**. Differences in ACS2 protein levels also support that *acs2-1* is an over-expression, and *acs2-2* is a reduced-expression mutant.

It is believed that ACS2 protein stability is regulated by protein phosphorylation at a conserved Ser-460 residue (Kamiyoshihara et al., 2010). Therefore, we examined whether mutation affected the phosphorylation status of ACS2 protein using a phospho-ser antibody. Considering that ACS2 protein level widely varied in *acs2-1*, WT, and *acs2-2*, the gel was run with 20 µg WT protein, 5 µg *acs2-1* protein, and 80 µg *acs2-2* protein, to have an equal amount of ACS2 protein in each lane. At equal levels of ACS2 protein loading, *acs2-1, acs2-2*, and WT lanes showed an equal amount of phosphorylated ACS2 protein, indicating that mutations did not affect ACS2 phosphorylation **(Figure 4G)**.

### *acs2* mutations alter phytohormones level in fruits

Analysis of mutant fruits revealed that *acs2* mutations also significantly influenced other plant hormones. Levels of zeatin, ABA, JA (Barring MG), MeJA, and SA were higher in *acs2-1* than WT. Contrastingly, the IAA level was lower at MG, and at later stages, it was similar to WT (**Figure 5A**). Notably, homozygous F_2_ *acs2-1*/*acs2-1* plants showed similar upregulation of zeatin, ABA, JA, MeJA, and SA compared to F_2_-WT (*ACS2/ACS2*). Contrasting to *acs2-1*, and compared to WT, levels of ABA, and SA was lower in *acs2-2* (Barring MG), and in homozygous F_2_ (*acs2-2*/*acs2-2*) plants. Levels of MeJA, JA (at RR), and zeatin (at MG, TUR) was also lower than WT in homozygous F_2_ (*acs2-2*/*acs2-2*) plants. IAA level in homozygous F_2_ (*acs2-2*/*acs2-2*) plants (In *acs2-2* only at TUR) was higher than WT **(Figure 5A)**. These results indicate the opposite influence of *acs2-1* and *acs2-2* on the levels of other hormones during ripening.

**Figure 5.**
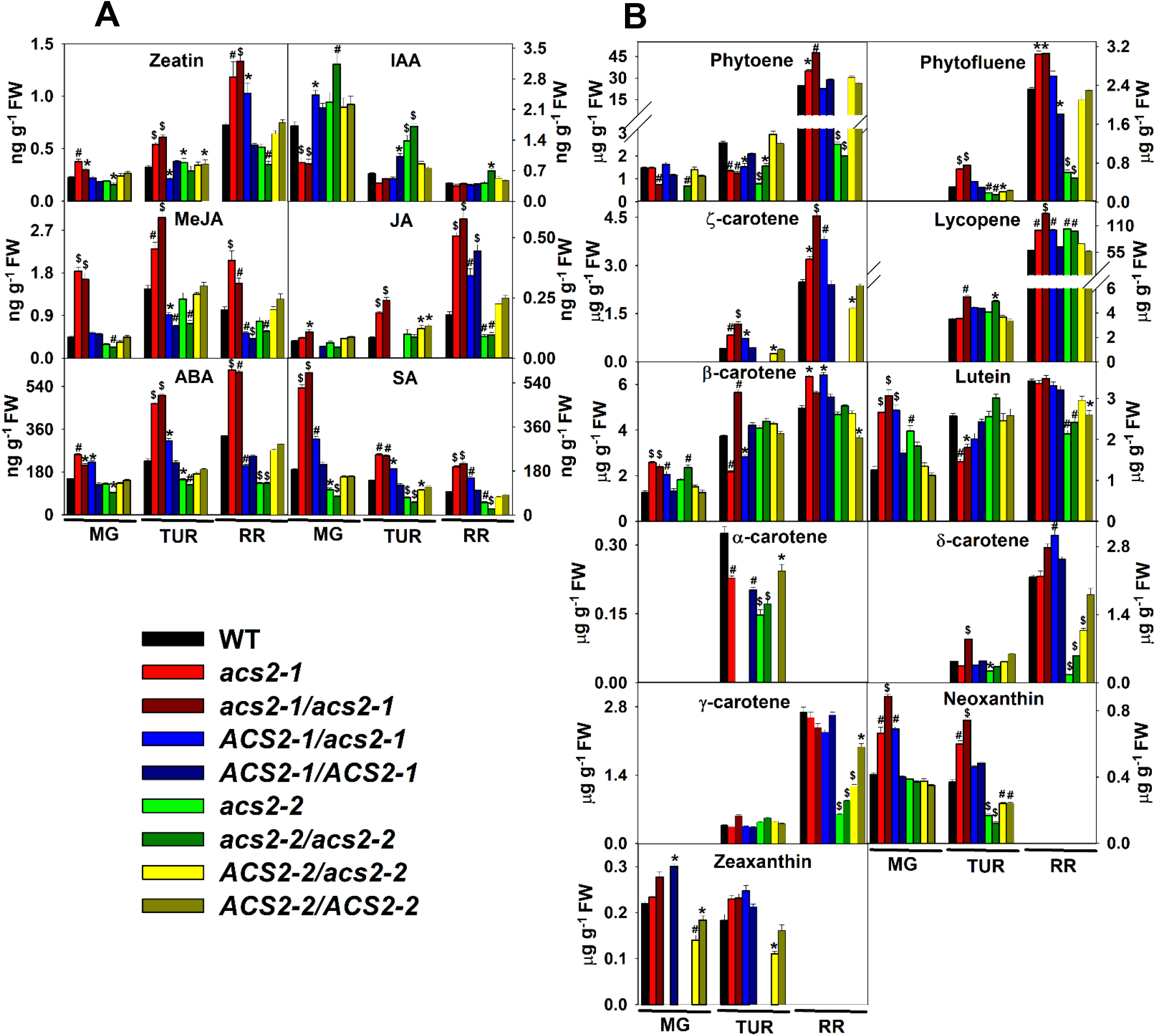
Phytohormones and carotenoids levels during ripening in WT, *acs2* mutants, and their BC_1_F_2_ progenies. (**A**) Phytohormones (**B**) Carotenoids. The levels of phytohormones were estimated using Liquid Chromatography-Mass Spectrometry (LC-MS). ABA (Abscisic acid), SA (Salicylic acid), IAA (Indole-3-acetic acid), JA (Jasmonic acid), MeJA (Methyl Jasmonic acid). The carotenoid levels were estimated using UPLC. The JA level in heterozygous (*ACS2*/*acs2–1*) at MG and TUR, and WT (*ACS2/ACS2*) F_2_ plants at TUR, were below the limit of detection. The BC_1_F_2_ progenies mentioned in this and following figures consisted of respective WT, heterozygous mutants, and homozygous mutants (WT-*ACS2–1/ACS2–1*; heterozygous-*ACS2–1*/*acs2–1;* homozygous mutant-*acs2–1*/*acs2–1*; WT-*ACS2–2/ACS2–2*; heterozygous-*ACS2–2*/*acs2–2;* homozygous mutant-*acs2–2*/*acs2–2*.)

### *acs2* mutants show higher carotenoid accumulation in fruits

The onset, progression, and completion of tomato ripening are visually monitored by the change of color imparted by carotenoids. While a total of 11 carotenoids were detected in fruits; only phytoene, lycopene, lutein, and β-carotene were detected at all ripening stages. In homozygous F_2_ (*acs2-1*/*acs2-1*) RR fruits, total carotenoid level was 1.85-fold higher than WT (*acs2-1*, 1.48-fold). The above increase was mainly contributed by a near doubling of lycopene and phytoene in *acs2-1*. Strikingly, *acs2-2* and homozygous F_2_ (*acs2-2*/*acs2-2*) RR fruits also had a higher total carotenoid level, mainly due to higher accumulation of lycopene. In *acs2-2*, levels of different carotenoids were distinctly different from both WT and *acs2-1* **(Figure 5B)**. The higher lycopene accumulation at the RR stage was also observed in BC_4_F_2_ plants of AV (*acs2-1*/*acs2-1*) and PED (*acs2-2/acs2-2*) substantiating the influence of respective *acs2* mutants on lycopene levels **(Supplemental Figure S4)**.

### *acs2* mutants oppositely affect metabolites level in fruits

Ripening of tomato is driven by extensive metabolic shifts making fruits palatable. Several of these changes are triggered by ethylene, as fruits of *Nr* mutant, which is deficient in ethylene perception, show different patterns of metabolic shifts than WT (Osorio et al., 2011). Consistent with the opposite nature of *acs2* mutants, PCA revealed distinct differences between metabolites of *acs2-2, acs2-1*, and WT **(Figure 6A, Supplemental dataset S4)**. Notably, PCA profiles of F_2_ fruits were distinctly closer to respective parents. PCA profile of *acs2-1* and homozygous F_2_ (*acs2-1*/*acs2-1*) overlapped. Similarly, PCA profiles of WT and homozygous F_2_ (*ACS2-1*/*ACS2-1*) distinctly overlapped at MG and TUR and were in close vicinity at RR. Strikingly, heterozygote F_2_ (*ACS2-1*/*acs2-1*) occupied an intermediate position in PCA between WT and *acs2-1* **(Figure 6B)**. A similar pattern in PCA profiles was also observed for *acs2-2*, where homozygous F_2_ mutant (*acs2-2*/*acs2-2*) overlapped with the respective parent, and heterozygous F_2_ (*ACS2-2*/*acs2-2*) showed intermediate PCA profile **(Figure 6C)**.

**Figure 6.**
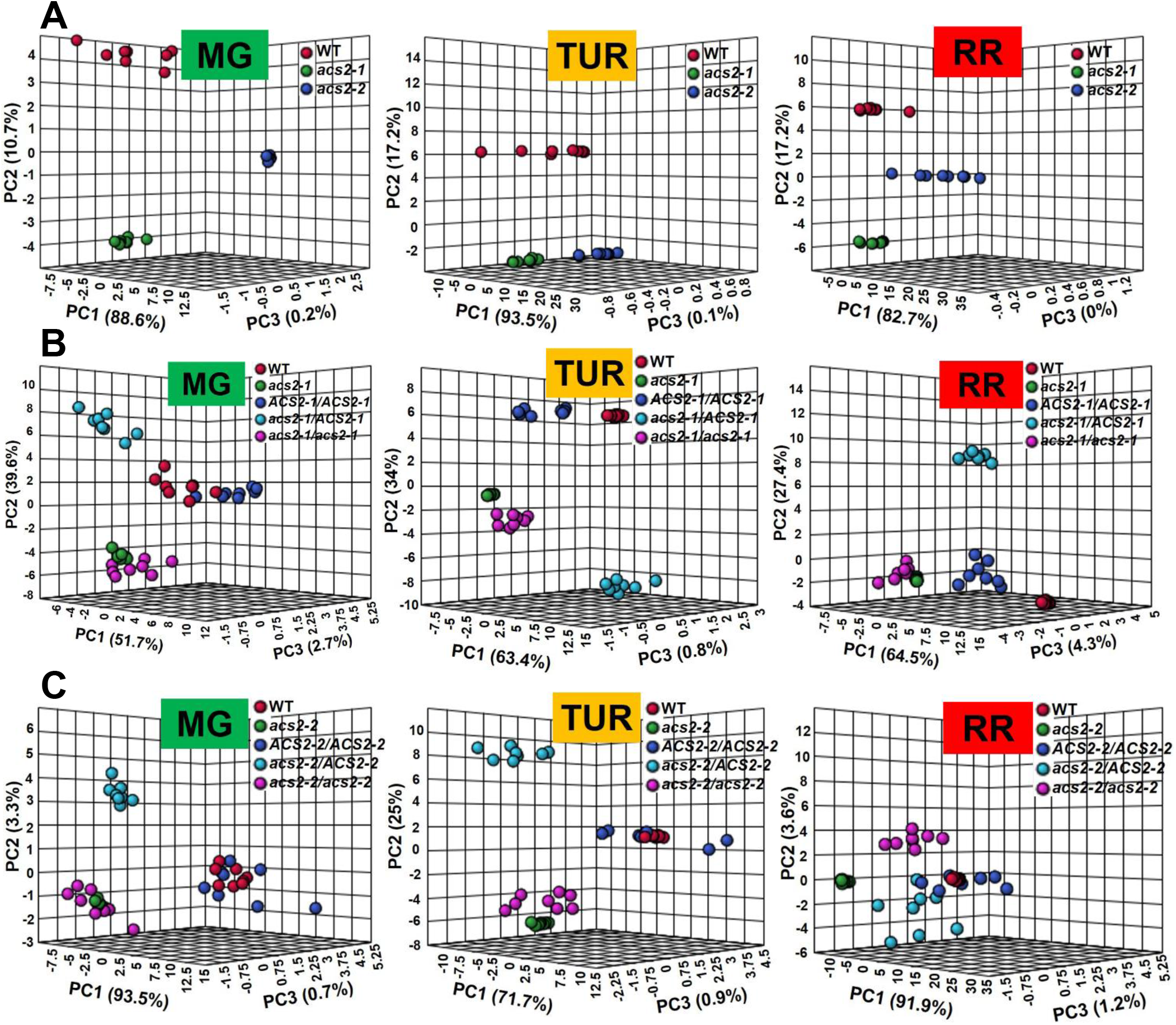
PCA of metabolites during ripening in WT, *asc2* mutants, and their BC_1_F_2_ progenies. **(A)** PCA of WT and *acs2* mutants. **(B)** PCA of WT, *acs2–1*, and its BC_1_F_2_ progeny. **(C)** PCA of WT*, acs2–2*, and its BC_1_F_2_ progeny.

Comparison of metabolite profiles of *acs2-1 and acs2-2* (**Figure 7**) and their backcrossed progeny with WT highlighted a strong linkage between the mutated gene and metabolome. The metabolome of *acs2-1* and F_2_ (*acs2-1/acs2-1)* was closer to each other than WT. Similarly, the metabolome of WT and F_2_ (*ACS2-1/ACS2-1)* showed only mild differences. The same was observed for metabolite profiles of *acs2-2* and F_2_ (*acs2-2/acs2-2)* fruits (**Figure 7, Supplemental dataset S4**). Metabolome of *ACS2-1/acs2-1* and *ACS2-2/acs2-2* was intermediate. Observed profiles of metabolome were also consistent with PCA profiles of mutated plants and their backcrossed progeny.

**Figure 7.**
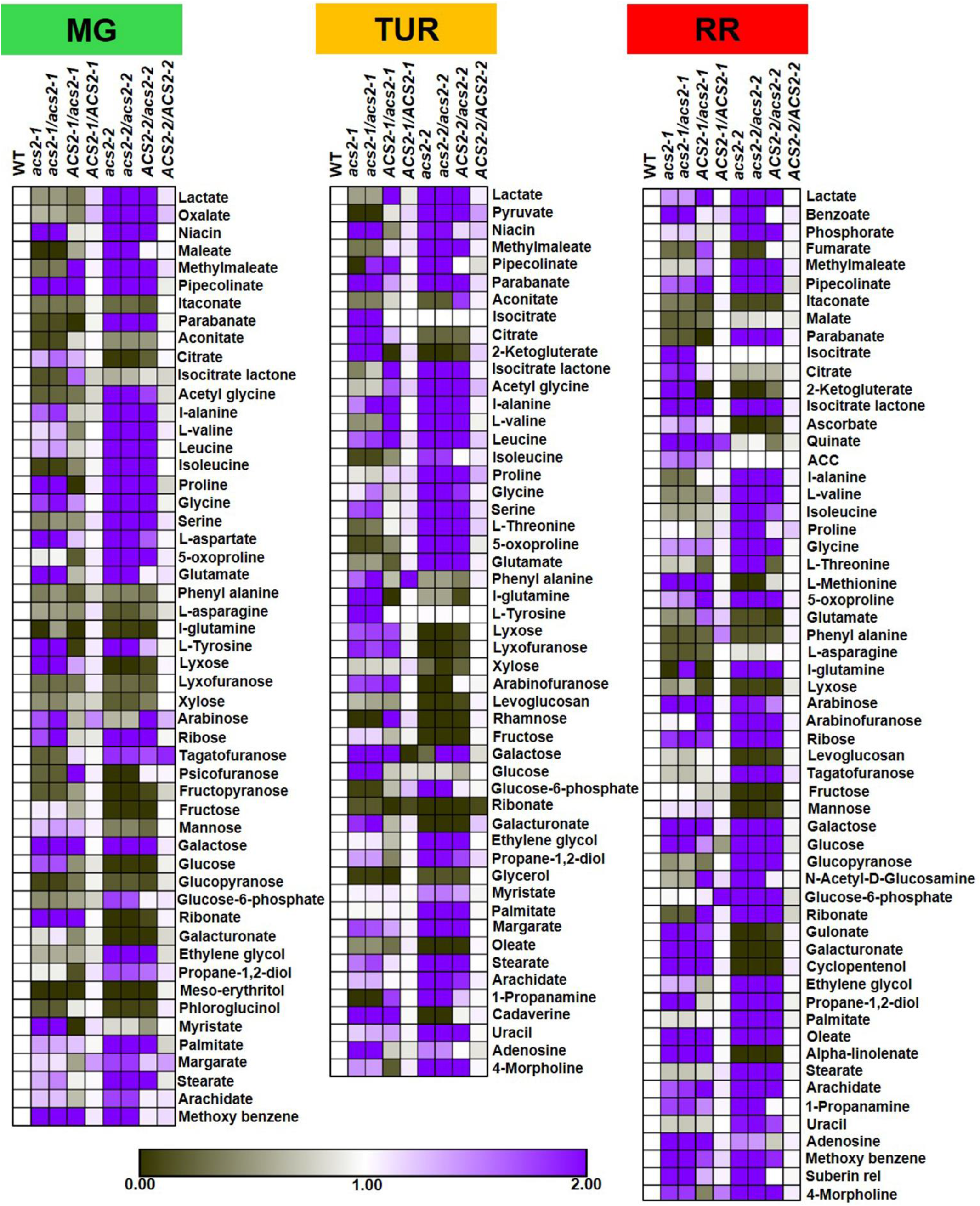
Relative levels of metabolites in ripening fruits of WT and mutants. Heat maps show differential expression of metabolites at different ripening stages in fruits of WT, *acs2–1*, *acs2–2*, and their respective BC_1_F_2_ progenies. The relative changes of metabolites represent the mutant/WT ratio, calculated using parental WT used for crosses, at respective ripening stages as the denominator. Only significantly up- or down-regulated metabolites are represented in the heat map (> 1.5 fold). For details of BC_1_F_2_ progenies, see Figure 5.

A comparison of metabolite profiles of *acs2-1* and *acs2-2* highlighted distinct differences between the two mutants. The effect of *acs2-1* on the relative up-/down-regulation of most metabolites was milder than *acs2-2*. In *acs2-2*, the majority of amino acids derived from the glycolysis pathway were upregulated, whereas *acs2-1* only mildly affected the aminome except for 10-fold higher methionine in RR fruits. Commensurate with high ethylene emission and methionine, ACC levels were also high in *acs2-1* fruits at the RR stage. In *acs2-2*, 2-oxoproline was upregulated at all ripening stages. Similar to aminome, most organic acids were downregulated in *acs2-1* fruits. Only a few organic acids were upregulated in *acs2-1*, such as citrate, α-ketoglutaric acid, isocitric acid (TUR, RR), and succinate (TUR). Similarly, only a few organic acids were upregulated in *acs2-2* [Lactate; Oxalate (MG, TUR); Phosphoric acid; Maleate (MG)]. Compared to *acs2-2*, fewer sugars were higher in *acs2-1* than WT. In *acs2-2*, sucrose and glucose-6-phosphate were higher than WT (**Supplemental figure 7**).

### *acs2-1* mutation elevates ETR3/4 expression

System II ethylene synthesis in ripening tomato fruits is associated with higher transcript levels of *ACS2*, *ACS4*, *ACO1, ACO2*, and *ACO3* (Liu et al., 2015). During ripening, transcript levels of *ACS2*, *ACS4*, *ACO1*, and *ACO2* were upregulated in *acs2-1*, in homozygote-F_2_ (*acs2-1*/*acs2-1*) and heterozygote-F_2_ (*ACS2-1*/*acs2-1*) at TUR and RR stages than parental WT and WT-F_2_ (*ACS2-1*/*ACS2-1*) (**Figure 8A**). Contrastingly, in *acs2-2*, in homozygote-F_2_ (*acs2-2*/*acs2-2*), and in heterozygote-F_2_ (*ACS2-2*/*acs2-2*) transcript levels of *ACS2* (barring RR in *acs2-2*), *ACS4* (barring TUR in *acs2-2*), *ACO1, ACO2* and *ACO4* (barring *ACS2-2*/*acs2-2*), were lower during ripening WT. Both *acs2-1* and *acs2-2* and their respective homozygous F_2_ progeny had no consistent influence on transcript levels of *ACO3*. Out of six *ETR* genes in tomato, *acs2-1* specifically upregulated *ETR3*, *ETR4* at the TUR stage **(Figure 8A)**. In contrast, *acs2-2* and its homozygote-F_2_ (*acs2-2*/*acs2-2*), downregulated *ETR1*, *ETR2*, and *ETR4* (barring *acs2-2* at BR) at all stages of ripening. For *ETR5*, only homozygote-F_2_ (*acs2-2*/*acs2-2*) showed downregulation at BR and RR stages. For *ETR6*, no consistent influence of *acs2*-*1* or *acs2*-*2* and their homozygote was observed.

**Figure 8.**
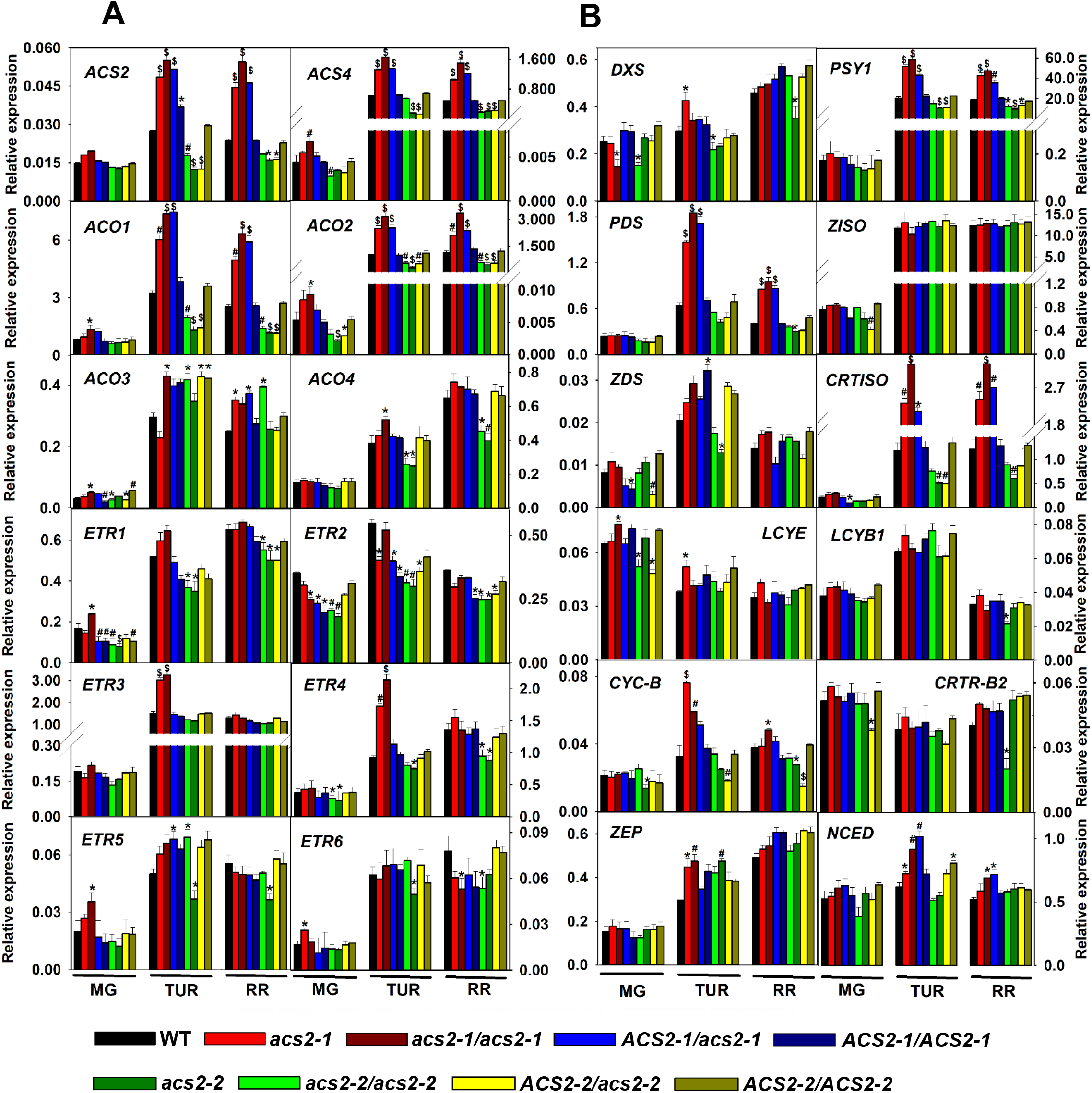
Relative expression of transcripts of ethylene receptors, ethylene and carotenoids biosynthesis genes during ripening in WT, *acs2* mutants, and their BC_1_F_2_ progenies. *ACS-1-aminocyclopropane carboxylic acid synthase; ACO-1-aminocyclopropane carboxylic acid oxidase; CRTISO*-*carotenoid* isomerase; *CRTR-B2-β-carotene hydroxylase 2; CYCB - chromoplast specific lycopene β-cyclase*; *DXS-deoxy-xylulose 5-phosphate synthase ETR-ethylene receptor; LCYB1*-*lycopene β-cyclase 1; LCYE-lycopene ε-cyclase; NCED-9-cis-epoxycarotenoid dioxygenase 1, PDS-phytoene desaturase*; *PSY1-phytoene synthase 1*; *ZDS*-*ζ-carotene desaturase; ZEP-zeaxanthin epoxidase; ZISO-ζ-carotene isomerase;* For BC_1_F_2_ progenies details, see Figure 5.

We next examined whether higher carotenoids level in *acs2-1* and *acs2-2* was also associated with increased expression of carotenoid biosynthesis genes. Influence of *acs2-1*, its homozygote-F_2_ (*acs2-1*/*acs2-1*), and heterozygote-F_2_ (*ACS2-1*/*acs2-1*) on the expression of carotenoid biosynthesis genes at TUR and RR stages was mainly confined to three genes of pathway-phytoene synthase1 (*PSY1*), phytoene desaturase (*PDS*), and carotenoid isomerase (*CRTISO*). Higher transcript levels of *PSY1*, *PDS*, and *CRTISO* in heterozygote-F_2_ (*ACS2-1*/*acs2-1*) fruits indicate a semi-dominant influence of *acs2-1* on these genes. *acs2-1* homozygote also upregulated chromoplast-specific lycopene-β-cyclase (*CYCB*). Besides the above genes, *acs2-1* upregulated *NCED1* and *ZEP* (only at TUR) expression. Interestingly, the expression of *PSY1* followed an opposing pattern in *acs2-2* with lower transcript levels. Few other genes in *acs2-2* were downregulated at specific stages, viz. *ZDS* (at TUR in homozygote), *CRITSO* (homo- and hetero-zygote at TUR, homozygote at RR), and *CYCB* (homo- and hetero-zygote at RR) **(Figure 8B)**.

## Discussion

Ethylene plays several roles throughout the life cycle of plants, regulating a plethora of developmental responses. Among these, ethylene regulation of the ripening of climacteric fruits has drawn the most attention. In this study, a comparison of two contrasting *acs2* mutants uncovers that besides regulating fruit ripening; ethylene produced from *ACS2* also modulates several aspects of tomato phenotype right from seed germination.

### Ethylene promotes seed germination

It is established that in higher plants, seed germination is antagonistically regulated by ABA and GA (Shu et al., 2016). Our results indicate that endogenous ethylene participates in tomato seed germination. Faster germination of *acs2-1* is likely related to higher ethylene emission. Slower germination of *acs2-2* associated with reduced ethylene emission conforms to the above assumption. Consistent with this, peak emission of ethylene from *acs2-1* seeds coincides with the completion of germination (Lashbrook et al., 1998). Considering that transgenic tomato seeds overexpressing *ERF2* show faster germination (Pirrello et al., 2006), it is likely that higher ethylene emission from *acs2-1* amplifies ethylene signaling.

Seedlings of *acs2-1* grown in the air do not manifest constitutive triple response alike Arabidopsis *ETO* mutants (Guzman and Ecker, 1990) or tomato *Epi* mutant (Barry et al., 2001). Since ethylene biosynthesis/action inhibitors do not revert *Epi* phenotype, *Epi* probably affects an ethylene-signaling component (Fujino et al., 1989; Barry et al., 2001). Considering that Arabidopsis *eto2* (*ACS5*) and *eto3* (*ACS9*) seedlings produce 20-90 fold higher ethylene than the wild-type, 3-4 fold higher ethylene emission by *acs2-1* seedlings may not be sufficient to elicit a constitutive triple response (Kieber et al., 1993; Chae et al., 2003). Nevertheless, higher ethylene emission from *acs2-1* seems to elicit shorter hypocotyls in etiolated seedlings than WT. Further shortening of hypocotyls in ethylene-treated seedlings likely reflects combined action of external and endogenous ethylene produced by seedlings, as hypocotyls shortening followed *acs2-1>*WT*>acs2-2*.

### Ethylene affects both vegetative and reproductive phenotypes

Contrasting phenotypes of *acs2-1* and *acs2-2* seem to result from high/low ethylene emission from respective mutants. Despite higher ethylene emission, *acs2-1* phenotype had no resemblance to *35S::ACS2* transgenic tomato displaying epinasty, reduced growth, and higher ethylene emission (Lee et al., 1997). Likewise, the *acs2-1* phenotype was also distinct from the tomato *Epi* mutant that displays an erect growth habit, curled leaves, thickened stems, and petioles. Notably, the *Epi* effect is restricted to vegetative development (Barry et al., 2001), whereas *acs2-1* affects both vegetative and reproductive phenotypes.

Contrasting phenotypes of *acs2-1* and *acs2-2*, while emanate from high/low ethylene emission, modulation of hormones and metabolites also underlie phenotypic differences. Hormonal profiles of *acs2-1* and *acs2-2* leaves indicated crosstalk between ethylene and other hormones. In conformity with reduced ethylene emission delays tomato leaf senescence (John et al., 1995), detached leaves of *acs2-1* showed faster senescence, while *acs2-2* showed slower senescence. In tomato plants grown under high salinity, onset and progression of leaf senescence correlated with endogenous zeatin/ACC ratio (Ghanem et al., 2008). Likewise, opposite zeatin and ethylene levels in *acs2-2* and *acs2-1* leaves imply that the ethylene/zeatin ratio is a key determinant regulating leaf senescence. Similarly, high ABA/ethylene level in *acs2-1* leaves agrees with a parallel increase in ABA and ethylene in salt-stressed tomato plants (Ghanem et al., 2008).

Generally, plants undergoing pathogen or insect damage show a concerted action of ethylene with SA and JA to activate defense responses (Yang et al., 2015). In tomato plants infected with Alternaria, ethylene and JA acted synergistically to promote susceptibility. Conversely, SA promoted resistance to Alternaria and antagonized ethylene signaling (Jia et al., 2013). Therefore, it is conceivable that higher ethylene emission in *acs2-1* affects levels of defense hormones such as JA and SA. The upregulation of JA, MeJA, and downregulation SA in *acs2-1* leaves, conforms with this assumption.

The relationship between auxin and ethylene is complex and has both synergistic and antagonistic interactions (Muday et al., 2012). Auxin and ethylene co-act to facilitate tomato root tip penetration in soil (Santisree et al., 2011). Conversely, ACC-treatment reduces free auxin in tomato roots indicating antagonistic action of ethylene (Negi et al., 2010). Reduction in IAA and IBA levels in *acs2-1* points toward an antagonistic action of ethylene on the IAA/IBA level. Since barring zeatin and SA, other hormones are not affected in *acs2-2* leaves; it seems that hormonal modulation in *acs2-1* leaves reflects a response analogous to stress, where on surpassing a threshold, ethylene modulates levels of stress-related hormones.

Distinct PCA profiles of *acs2* mutants are consistent with their contrasting responses. The cross-comparison showed that most metabolites were downregulated in *acs2-1*, while in *acs2-2*, several were upregulated than WT. Ostensibly, altered ethylene emission from *acs2-1* and *acs2-2* leaves affects metabolites profiles oppositely. Higher citrate, isocitrate levels, and lower malate levels in *acs2-1* leaves indicate an upsurge in respiratory metabolism, a hallmark of stress. Purportedly, citrate accumulation is construed as a response mechanism to alleviate stress (Gupta et al., 2012). Differences in ethylene emission also influence the reproductive phenotype, as both mutants display opposite flower numbers and fruit set than WT. Taken together, it is implicit that distinct phytohormone and metabolite profiles of *acs2* mutants underlie differences in their phenotypes.

### *acs2* mutants show contrasting ripening progression

It is believed that *ACS2* contributes to system-II ethylene emission, whereas its role in system-I ethylene emission is minimal (Barry et al., 2000). Nonetheless, faster progression to the MG stage in *acs2-1* implies a role for *ACS2* in system-I ethylene emission. Consistent with this, ethylene-treated Micro-Tom fruits, too showed faster attainment of the MG stage (Kevany et al., 2007). Our results support the view that the transition from MG to RR stage is linked with ethylene-induced ripening. Post-MG stage, faster progression to RR stage, and the onset of senescence in *acs2-1* are consistent with the pivotal role of ACS2 in the progression of ripening. The above assumption is also corroborated with slower progression to the RR stage and delayed senescence in *acs2-2* fruits. Shorter MG to RR transition period in F_2_-*acs2-1/acs2-1* and a far-longer period in F_2_-*acs2-2/acs2-2* progeny of M82, AV and PED, further corroborate close linkage with ACS2.

### Ethylene also influences the expression of *ETR3/4* genes

The higher transcript level of *ETR3* and *ETR4* in *acs2-1* fruits at the TUR stage indicates the likelihood of stimulation by ethylene (Lashbrook et al., 1998; Kevany et al., 2007). Lowered transcript levels of *ETR1, 2, 4, 5* in *acs2-2* indicates that their expression is also related to ethylene levels (Okabe et al., 2011; Mubarok et al., 2019). Higher *ACS2*, *ACS4*, *ACO1*, and *ACO2* transcript levels in ripening *acs2-1* fruits conform with the notion that ethylene in an autocatalytic fashion enhances its synthesis. Consistent with this, reduced ethylene emission from *acs2-2* fruits lowers *ACS2*, *ACS4*, *ACO1, ACO2*, and *ACO4* transcript levels. Taken together, it appears that higher ethylene emission from *acs2-1* fruits promotes higher transcript levels of ripening-specific ethylene biosynthesis genes and ethylene receptors, and it is opposite in *acs2-2*.

### ACS activity of mutants is correlated with ACS2 protein levels

The *in vitro* assays corroborated that higher ethylene emission was causally related to ACS activity. In ripening tomato fruits, endogenous ACC content closely correlates with ethylene emission (Hoffman and Yang, 1980). Higher ethylene emission from *acs2-1* fruits also correlates with higher ACC as well as MACC levels. Also, *acs2-1* fruits had higher ACS and ACO enzyme activities. Considering that *acs2-1* fruits accumulate 4-fold higher ACS2 protein, it may have contributed to higher ACS activity. Conversely, lower ethylene emission from *acs2-2* was associated with lower ACC and MACC levels, reduced ACS, and ACO enzyme activities and lower amounts of ACS2 protein.

Substantial differences in immuno-detectable ACS2 protein in *acs2-1* and *acs2-2* fruits indicated that the above mutations likely affect transcription or stability of the protein. The likelihood of enhanced stability of *ACS2* protein in *acs2-1* by post-translational modification seems to be remote. Typical of type-I ACS, tomato ACS2 protein has three conserved serine phosphorylation sites in C-terminal (Kamiyoshihara et al., 2010). However, there was no difference in phosphorylation magnitude in ACS2 protein in mutants and WT. In Arabidopsis *eto2* (ACS5) and *eto3* (ACS9) mutants encoding Type-II ACS, mutations close to C-terminal stabilizes ACS protein by making it resistant to E3 ligase dependent proteolysis (Chae and Kieber, 2005). However, Type-I ACS proteins are not subjected to ubiquitin-mediated proteolysis. It remains possible that α-helix change by V352E confers protein stability or stimulates ACS2 catalytic activity. The above possibility is supported by *in silico* analysis showing higher stability for ACS2-1 protein than WT, which may have contributed to increased ACS2 levels in *acs2-1*.

Notwithstanding predicted higher ACS2-1 protein stability, increased *ACS2* transcript level in *acs2-1* seems to also contribute to the higher amount of ACS2 protein. The abundance of mRNA is governed by a combination of transcription, splicing, and turnover. A398G mutation in *acs2-1* is located at the 5′ splice site of second-exon and third-intron junction. The presence of G at splice junction is predicted to confer more efficient mRNA splicing than WT (Hebsgaard et al., 1996), which in turn may boost *ACS2* transcript levels. A large-scale analysis revealed that synonymous alleles bearing guanine or cytosine at the third position of a codon increased mRNA half-life (Duan et al., 2013). Considering A398G (K=) is a synonymous mutation, it may have enhanced half-life of *acs2-1* mRNA. During the climacteric phase of tomato ripening, ethylene upregulates *ACS2* transcripts in a positive feedback fashion (Alba et al., 2005). It is plausible that higher ethylene emission from *acs2-1* fruits in an autocatalytic fashion boosted *ACS2* transcript levels, which in turn led to a higher level of ACS2 protein. Available evidence points that enhanced transcript levels of *acs2-1* and a likely increase in protein stability seem to contribute to the increased ACS2 protein level.

Considering *acs2-2* harbors two mutations in the promoter, the lower level of ACS2 protein seems to be a consequence of reduced *ACS2* transcript level in *acs2-2* fruits. These mutations most likely affect transcript levels of *ACS2* by perturbing interactions between transcription factors and their binding sites. Among transcription factors interacting with mutated sites, AZF is a known repressor of ABA-mediated gene expression (Kodaira et al., 2011). It is plausible that the gain of the AZF binding site due to T-106A mutation in *the acs2-2* promoter may reduce its expression. Besides, mutation at T-382A may hinder demethylation during ripening, thus affecting *acs2-2* transcript levels. Anyhow, contrasting transcript, ACS2 protein levels, ACS enzyme activity, and ethylene emission support the notion that *acs2-1* is a hypomorphic enhanced-expression, and *acs2-2* is a reduced-expression mutation.

### *acs2* mutants also influence other phytohormones in fruits

While ethylene is considered a master regulator of tomato ripening, it may crosstalk and influence other hormones during ripening. Consistent with the above, hormone profiling revealed alteration in other phytohormones’ levels in *acs2* mutants. *acs2-1* displayed higher levels of zeatin, ABA, JA, MeJA, and SA. Downregulation of ABA and SA in *acs2-2* conforms to its opposite action. Higher auxin levels in *acs2-2/acs2-2* (MG, TUR) fruits are consistent with reported auxin/ethylene antagonism during the early phase of tomato ripening (Su et al., 2015). It is reported that ABA acts as a positive regulator of ethylene biosynthesis at the onset of ripening in tomato (Zhang et al., 2009). Our results indicate that ethylene acts as a positive regulator of ABA in ripening fruits, as ABA levels are higher in *acs2-1* and lower in *acs2-2*. Increase in the ABA level is probably mediated by the upregulation of *NCED1* expression in *acs2-1*, a gene involved in ABA biosynthesis. Considering that both JA and MeJa levels are higher in *acs2-1* and MeJa levels are lower in *acs2-2/acs-2-2*, there seems to be a positive correlation between ethylene and JA levels in tomato fruits. In conformity with this, jasmonate-deficient tomato mutants show lower ethylene emission during fruit ripening (Liu et al., 2012). Interaction of ethylene with phytohormones also seems to be organ-specific, as evidenced by lower zeatin levels in *acs2-1* leaf, whereas fruits have higher zeatin levels. Whilst ethylene modulates hormonal level in ripening fruits, the interrelationship among these hormones is complex, and likely involves the intersection of various regulatory pathways (Breitel et al., 2016).

### *ACS2* mutants show high lycopene level in fruits

The onset of fruit-specific carotenogenesis is closely linked with ethylene biosynthesis as antisense suppression of *ACS2* (Theologis et al., 1993) or defect in ethylene perception in *Nr* mutant (Lanahan et al., 1994) inhibits characteristic red coloration of tomato fruits. Higher carotenoids level in *acs2-1* is consistent with the linkage between ethylene emission and carotenoid accumulation. Analogous to influence on the hormonal level, *acs2-1* in a semi-dominant fashion boosted carotenoid levels in heterozygous *acs2-1* plants. Ethylene seems to stimulate carotenoid levels in *acs2-1* by upregulating transcripts of *PSY1*, *PDS*, and *CRTISO*, the key genes regulating phytoene and lycopene biosynthesis. Surprisingly, though most carotenoid intermediates and key genes transcripts such a *PSY1* and *CRTISO* were downregulated in *acs2-2*, yet it accumulated a higher level of lycopene. In tomato fruits, carotenoid biosynthesis is geared towards lycopene accumulation, which is sequestered and stored in plastoglobules and lycopene crystals in chromoplasts (Kilambi et al., 2013; Nogueira et al., 2013). It is likely that prolonged transition period from MG to RR stage in *acs2-2* facilitated continued synthesis and sequestration of lycopene, in turn leading to higher lycopene levels despite having reduced transcript levels of carotenogenic genes. This view is also substantiated by higher lycopene accumulation in the introgressed progeny of *acs2-1, acs2-2* in AV, and PED cultivars.

### *ACS2* mutants show contrasting effects on metabolite profiles

Consistent with the pleiotropic effect of ethylene, levels of several metabolites in fruits were altered in *acs2-1* and *acs2-2*. Different PCA profiles of *acs2-1, acs2-2*, and WT are consistent with the view that hormonal signaling perturbation strongly influences metabolite levels (Bastías et al., 2014). Though *acs2-2* and *Nr* mutant are genetically distinct, a comparison of *acs2-2* and *Nr* metabolite profiles (Osorio et al., 2011) showed a similar shift in levels of at least 13 metabolites from MG to RR stage including key TCA cycle constituents such as citric acid, isocitric acid, and malic acid. This entails that the above similarity in metabolites between *Nr* and *acs2-2* likely emanates from reduced ethylene-mediated signal transduction in respective mutants.

Reduced CO_2_ emission from *acs2-2* fruits seems to be linked with reduced ethylene emission (Tigchelaar et al., 1978). Higher levels of amino acids in *acs2-2* may be linked with reduced respiration diverting glycolysis/TCA cycle intermediates to amino acid synthesis. Likewise, reduced ACS2 activity in *acs2-2* likely leads to high 5-oxoproline, which is part of the reactions needed to close the methionine salvage cycle in plants (Ellens et al., 2015). Low levels of 2-ketoglutarate in *acs2-2*, where 5-oxoproline accumulates, and high level in *acs2-1*, where 5-oxoproline does not accumulate, indicates that methionine salvage cycle differently operates in *acs2-1* and *acs2-2*, to prevent the accumulation of α-ketoglutaramate, a toxic molecule (Cooper, 2004).

Our study highlights that perturbation in the function of a single gene distinctly influences a wide range of metabolites. Importantly, metabolic shifts associated with *acs2-2* and *acs2-1* is largely retained in their backcrossed homozygous progeny. Likewise, in backcrossed homozygous WT, metabolic profiles revert to parental WT. Studies on the genetic control of metabolism have underscored QTLs as major determinants of metabolic shifts and profiles (Fernie and Tohge, 2017). Co-segregation of metabolite profiles with respective *acs2* mutations conforms with the above view. Importantly, it also highlights that an increase/decrease in ethylene levels in mutants has a cascading effect on plant metabolism, which in turn, accelerates/decelerates the ripening process.

In conclusion, our results highlight that besides a pivotal role in fruit ripening, *ACS2* participates in several facets of tomato development. Faster germination of *acs2-1* implies the role of *ACS2* in seed germination. Contrasting vegetative and reproductive phenotypes of mutants, including leaf/fruit senescence, indicate a role of *ACS2*, as a key ethylene biosynthesis enzyme, during tomato development. Faster transition to the MG stage in *acs2-1* mutant suggests the role of *ACS2* in early fruit development. Our study also entails that alleles of *ACS2* can be potentially used for modulating on-vine tomato ripening.

## Materials and Methods

### Mutant isolation

Two EMS-mutagenized populations of tomato (*Solanum lycopersicum*) cultivars, M82 (Menda et al., 2004), and Arka Vikas were used for screening of mutants. Genomic DNA was isolated from 8-fold pooled plants (Sreelakshmi et al., 2010), and mutation detection was carried out on the LICOR-4300 DNA analyzer (TILL et al., 2006) (**Supplemental Table 9)**. The presence of mutated gene copy and its zygosity in backcrossed plants AV, M82, and PED was monitored by the CEL-I endonuclease assay (Mohan et al., 2016).

### Characterization of mutants

The seeds sterilized with 4% (v/v) NaOCl were sown on filter papers, and germination was visually monitored. For phenotype studies, after sowing on agar (0.8% w/v), seedlings were grown in light (100 µmol/m^2^/s) or darkness. For ethylene-mediated growth inhibition, seedlings were grown in darkness in airtight plastic boxes, and at the onset of the experiment, a known volume of ethylene was injected in the box.

Plants were grown in the greenhouse under natural photoperiod (12-14h day, 28±1°C; 10-12h night, 14-18°C). For senescence, pigments, hormonal levels, and ethylene emission studies, leaves were harvested from the seventh node of 45-day-old plants. The floral and inflorescence morphology were from the second and third truss. For senescence study, the leaves laid on moist filter papers in Petri plates were placed under white light (100 µmol/m^2^/s), or darkness. The fruit development was monitored on-vine from the day of post-anthesis through different ripening stages until the fruit skin wrinkled. The firmness of detached fruits, pH, and °Brix was measured as described in Gupta et al. (2014).

### Estimation of ethylene and CO_2_ emission

The ethylene emission was monitored using a previously described procedure (Kilambi et al., 2013). The seedlings were transferred to an airtight glass vial on 0.8% (w/v) agar for 24h. For leaves, the detached leaves were placed on moistened filter paper and enclosed in airtight Petri plates for 4h. For ethylene emission, WT and mutant fruits were harvested from the second truss. The fruits were placed in an airtight container for 4h. At the end of the above-mentioned periods, one mL of headspace was withdrawn from the respective containers to estimate ethylene. For CO_2_ emission, the fruits were enclosed in a chamber containing the CO_2_ sensor (Vernier, OR, USA) for 10 min. After that, the CO_2_ emission was monitored using preinstalled Graphical Analysis^TM^ software.

### Estimation of ACC levels, ACS and ACO activity

The ACC, MACC levels, and ACS, ACO activity, were estimated using closed vials and calculated using Bulens et al. (2011) protocol. For ACC estimation, the fruit tissues were homogenized in 5% (w/v) sulphosalicylic acid, followed by 10 mM HgCl_2_ addition and release of ethylene by addition of 4% (w/v) NaOCl and saturated NaOH (2:1, v/v). To determine ACS (1gm tissue) and ACO (500mg tissue) activity, fruit tissue was homogenized in liquid N_2_ and mixed with the extraction buffer by vortexing. The supernatant obtained after centrifugation (21,000*g*) was desalted on the Sephadex-G25 column. The eluted fraction was assayed for ACS activity in a reaction mixture containing S-adenosyl-L-methionine. The reaction was terminated by adding HgCl_2,_ and ethylene was released by adding NaOCl/NaOH mixture. The ACO activity was estimated in a reaction mixture containing 1 mM ACC. For all of the above, at the end of the reaction, one mL headspace was withdrawn for ethylene estimation.

### Western blotting

The polyclonal antibodies were raised in rabbits against ACS2 protein using a peptide “EHGENSPYFDGWKAYDSD” custom synthesized by PEPTIDE-2.0, USA. The Ig fraction was purified using DEAE-Sepharose column using standard protocols. The fruits were homogenized in Tricine buffer (200 mM, pH 8.5), 2 mM pyridoxal-5-phosphate, 10 mM DTT with 50 mg PVPP. The homogenate was centrifuged (21,000*g*, 4°C) and the supernatant was desalted on Sephadex G25 column. Given the very low abundance of ACS2 protein, the eluted sample was immunoprecipitated after protein estimation (Bradford, 1976). The supernatant was gently shaken with purified IgG fraction at 37°C for 1h, followed by 1h shaking with protein A-Sepharose at 4°C. The beads were recovered by centrifugation (12,000g, 4°C). The supernatants were analyzed for the reduction in ACS2 activity, and protein A-Sepharose beads were used for Western blotting.

The immunoprecipitated ACS2 protein was recovered by boiling in SDS-PAGE loading buffer and separated in 12% gel following Laemmli (1970) protocol. The gel was electroblotted on the PVDF membrane, followed by Western blotting (Towbin et al., 1979). The non-specific binding sites on the membrane were blocked by incubating with milk powder. The membrane was first incubated with ACS2-antibody (1/1500 dilution), followed by incubation with an anti-rabbit IgG goat antibody (1/80,000 dilution), coupled with alkaline phosphatase. The alkaline phosphatase activity was visualized using standard BCIP-NBT assay.

### Profiling of phytohormones, carotenoids, and metabolites

The phytohormone levels were determined from leaves from the 7^th^ node of 45-day-old plants and fruits using Orbitrap Exactive-plus LC-MS following the protocol described earlier (Pan et al., 2010; Bodanapu et al., 2016). Carotenoid profiling was carried out following the procedure of Gupta et al. (2015). Metabolite analysis by GC-MS was carried out by a method modified from Roessner et al. (2000) described in Bodanapu et al. (2016).

### Real-time PCR

The RNA was isolated from fruit pericarp using TRI reagent (Sigma), and 2 µg DNase-treated RNA was reverse transcribed using a cDNA Synthesis Kit (Agilent), following the respective manufacturer’s protocol. RT-PCR was performed using iTaq Universal SYBR Green Supermix (BioRad) (**Supplemental Table S10)**. The relative differences were determined by normalizing Ct values of each gene to the mean expression of *β-Actin* and *Ubiquitin* genes and calculated using the equation 2^−∆Ct^.

### *In silico* characterization

The target protein structure and ACS2 sequence (PDB ID: 1IAX) (Huai et al., 2001) (https://www.rcsb.org/structure/1IAX), having 2.8Å resolution was retrieved from protein data bank (http://www.rcsb.org/pdb/). The 3-dimensional structure of *ACS2-1* (V352E) mutant and WT proteins were visualized using PyMol (http://www.pymol.org). The stabilization of *ACS2-1* versus WT protein was analyzed using the following software CUPSAT (http://cupsat.tu-bs.de/), MAESTROWeb (https://pbwww.che.sbg.ac.at/maestro/web), PoPMuSiCv3.1 (https://soft.dezyme.com/query/create/pop), STRUM (https://zhanglab.ccmb.med.umich.edu/STRUM/), and DynaMut (http://biosig.unimelb.edu.au/dynamut/). Promoter analysis was performed using PCbase (http://pcbase.itps.ncku.edu.tw/) and manually for ripening-specific transcription factors.

### Statistical analysis

All results are expressed as mean±SE of three or more independent replicates. The StatisticalAnalysisOnMicrosoft-Excel (http://prime.psc.riken.jp-/MetabolomicsSoftware/StatisticalAnalysisOn-MicrosoftExcel/) was used to obtain significant differences between data points. Heat maps and 3D-PCA plots were generated using Morpheus (https://software.broadinstitute.org/morpheus/) and MetaboAnalyst-4.0 (https://www.metaboanalyst.ca/), respectively. The statistical significance was determined using Student’s t-test (* for P≤0.05, # for P≤0.01 and $ for P≤0.001).

## Supplemental Data

**Supplemental Figure S1.** Splice site analysis of *acs2-1* mutant by NetGene2 server.

**Supplemental Figure S2.** Chlorophyll, carotenoids, and xanthophylls levels in the leaf of WT, *acs2-1*, and *acs2-2* mutant.

**Supplemental Figure S3.** Fruit firmness, total soluble solids (Brix) and pH of WT, *acs2* mutants, and their BC_1_F_2_ progenies at different ripening stages

**Supplemental Figure S4.** On-vine ripening, ethylene emission, and lycopene levels in BC_4_F_2_ progeny (*acs2-1*/*acs2-1)*, (*acs2-2*/*acs2-2*) in AV and PED cultivars.

**Supplemental Figure S5.** Raising and validation of antibodies specific for ACS2 peptides.

**Supplemental Figure S6.** Immunoprecipitation and Western blotting of ACS2 protein using purified IgG fraction.

**Supplemental Figure S7.** The metabolic shifts in *acs2* mutant fruits during ripening in comparison to WT.

**Supplemental Table S1.** EMS-mutagenized tomato populations used for the isolation of *ACS2* mutants.

**Supplemental Table S2.** *ACS2* mutant lines identified and confirmed for mutation.

**Supplemental Table S3.** The genetic segregation of *acs2-1* and *acs2-2* mutants in BC_1_F_2_ generation.

**Supplemental Table S4.** Increases in ACS2-1 protein stability predicted by different software.

**Supplemental Table S5**. The alteration of bonding pattern in ACS2 protein in *acs2-1* mutant compared to WT protein.

**Supplemental Table S6.** Disruption in transcription factor binding site in *acs2-2* due to promoter mutation.

**Supplemental Table S7.** Changes in the methylation status of *ACS2* promoter at −106 and −382 position during fruit ripening.

**Supplemental Table S8.** Comparisons of flower numbers and fruit set in WT and *acs2* mutants.

**Supplemental Table S9.** List of primers used for screening for mutations in *ACS2* by TILLING

**Supplemental Table S10.** List of genes and the primers used for qRT-PCR analysis.

**Supplemental dataset S1.** The transcription factor binding sites in the *ACS2* and *acs2-2* promoter predicted by PCbase.

**Supplemental dataset S2.** Changes in the methylation status of *ACS2* promoter during fruit ripening.

**Supplemental dataset S3.** List of metabolites identified in leaves of WT and *acs2* mutants.

**Supplemental dataset S4.** List of metabolites identified at different fruit ripening stages of WT and *acs2* mutants.

## ACKNOWLEDGMENTS

We thank Dr. Dani Zamir for EMS-mutagenized M82 cultivar lines and Dr. Alok Sinha for Phospho-Ser antibody. S.G. was recipient of DST Young Scientist grant and K.S. was recipient of UGC-JRF. S.S. gratefully acknowledges the financial support from the DBT-RA Program in Biotechnology and Life Sciences.

---

**Supplemental Figure S1.**
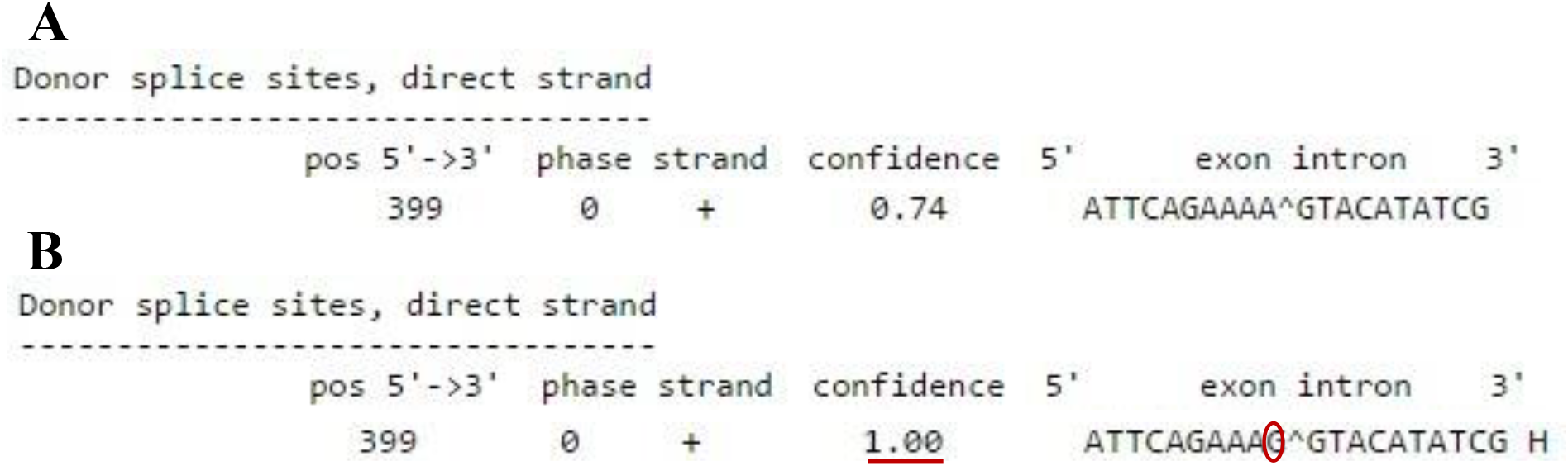
Splice site analysis of *acs2–1* mutant by NetGene2 server. **A** represents the WT splice site, while **B** represents the mutant splice site. Red circle denotes the site of changed base in mutant. The increase in confidence at 5′ denotes more efficient splicing of transcript.

**Supplemental Figure S2.**
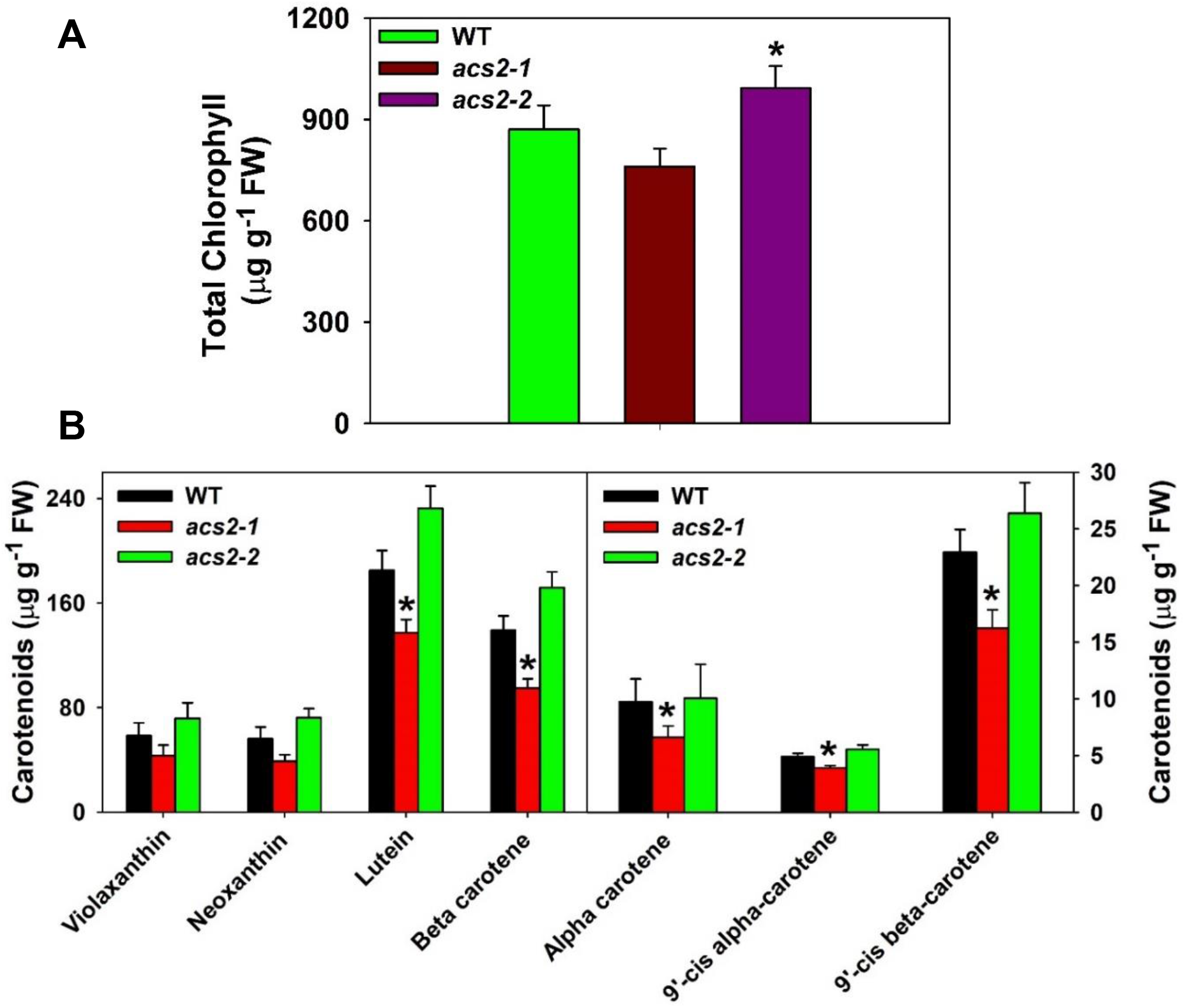
Chlorophyll (**A**), carotenoids, and xanthophyll (**B**) levels in leaf tissue of WT, *acs2–1*, and *acs2–2* mutant. Leaves were harvested from the seventh node of 45-days-old plants of wild type and mutants. (Student’s t-test; * for P ≤ 0.05, # for P ≤ 0.01 and $ for P ≤ 0.001, for chlorophyll and carotenoid estimation, sample number, n= 5 ± SE).

**Supplemental Figure S3.**
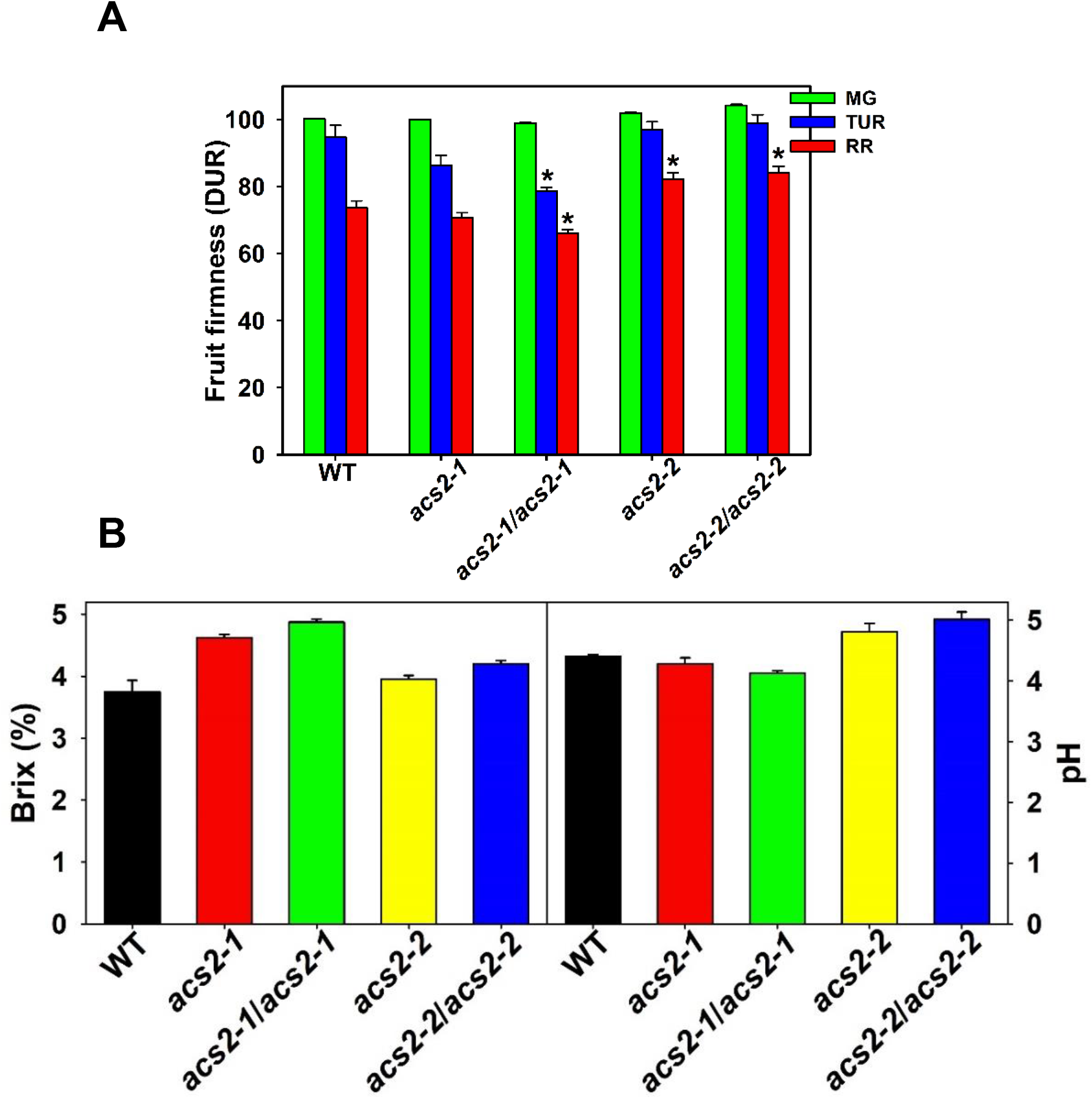
Fruit firmness (**A**), total soluble solids (Brix) (**B**), and pH (**C**) of WT, *acs2* mutants, and their BC_1_F_2_ progenies at different ripening stages. Firmness value was recorded by measuring each fruit at equatorial plane two-three times. (Student’s t-test; * for P ≤ 0.05, # for P ≤ 0.01 and $ for P ≤ 0.001, n=3 ± SE).

**Supplemental Figure S4.**
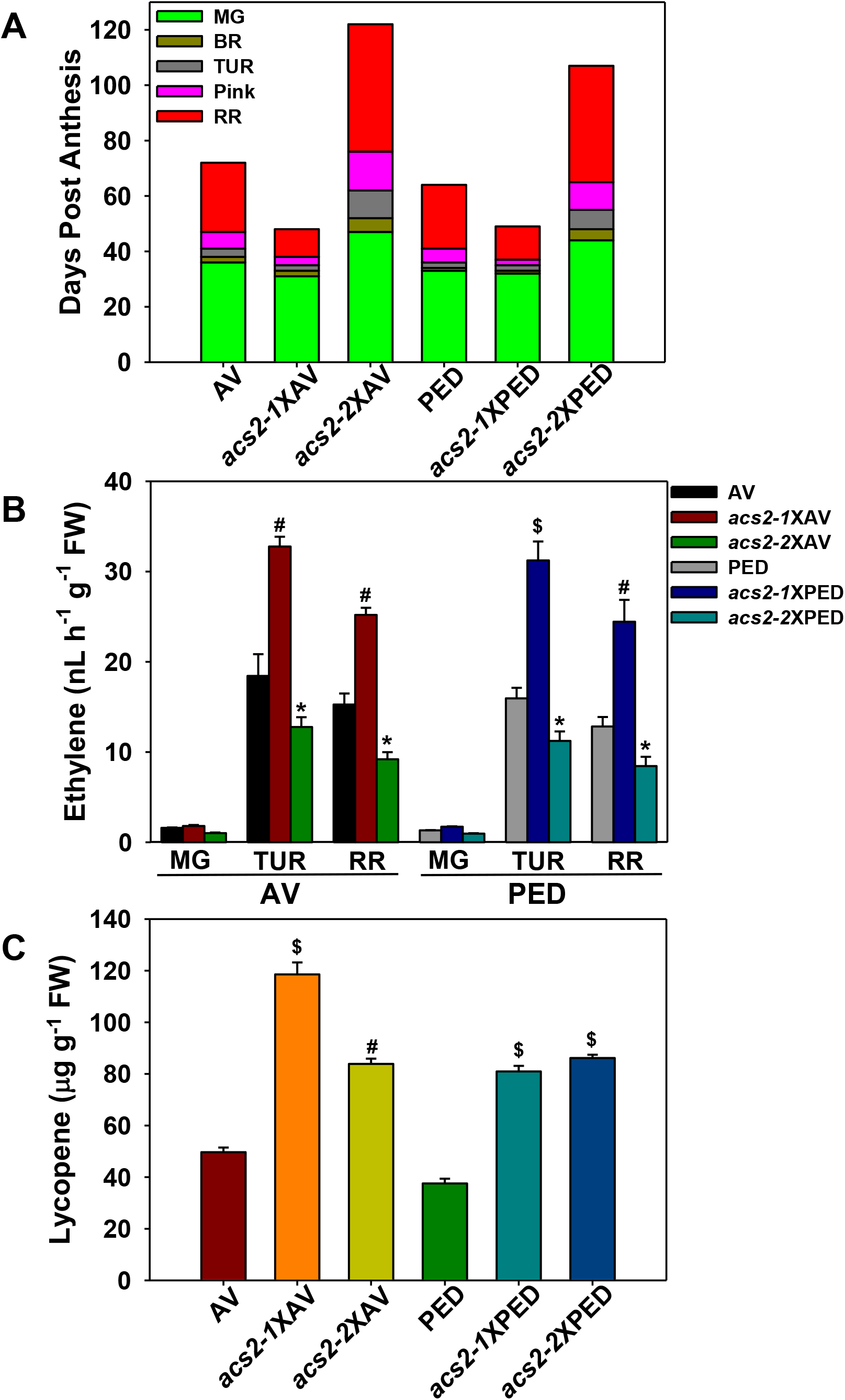
On-vine ripening, ethylene emission, and lycopene levels in BC_4_F_2_ progeny (*acs2–1*/*acs2–1)* (*acs2–2*/*acs2–2*) of Arka Vikas (AV), and Pusa Early Dwarf (PED). (**A**) Comparison of on-vine ripening (**B**) Ethylene emission during fruit ripening (**C**) Lycopene levels in red ripe fruits. Characterization was done in BC_4_F_2_ introgressed lines homozygous for respective mutations (*acs2–1* X AV= BC_4_F_2_ *acs2–1/acs2–1; acs2–2* X AV= BC_4_F_2_ *acs2–2/acs2–2; acs2–1* X PED*=* BC_4_F_2_ *acs2–1/acs2–1; acs2–1* X PED= BC_4_F_2_ *acs2–2/acs2–2*). **Note**, the introgressed lines show retention of fast/slow ripening, high/low ethylene emission, and high lycopene levels in red ripe fruits similar to *acs2–1* and *acs2–2* mutant lines backcrossed in M82 cultivar. The statistical significance was determined using Student’s t-test. (* for P ≤ 0.05, # for P ≤ 0.01 and $ for P ≤ 0.001).

**Supplemental Figure S5.**
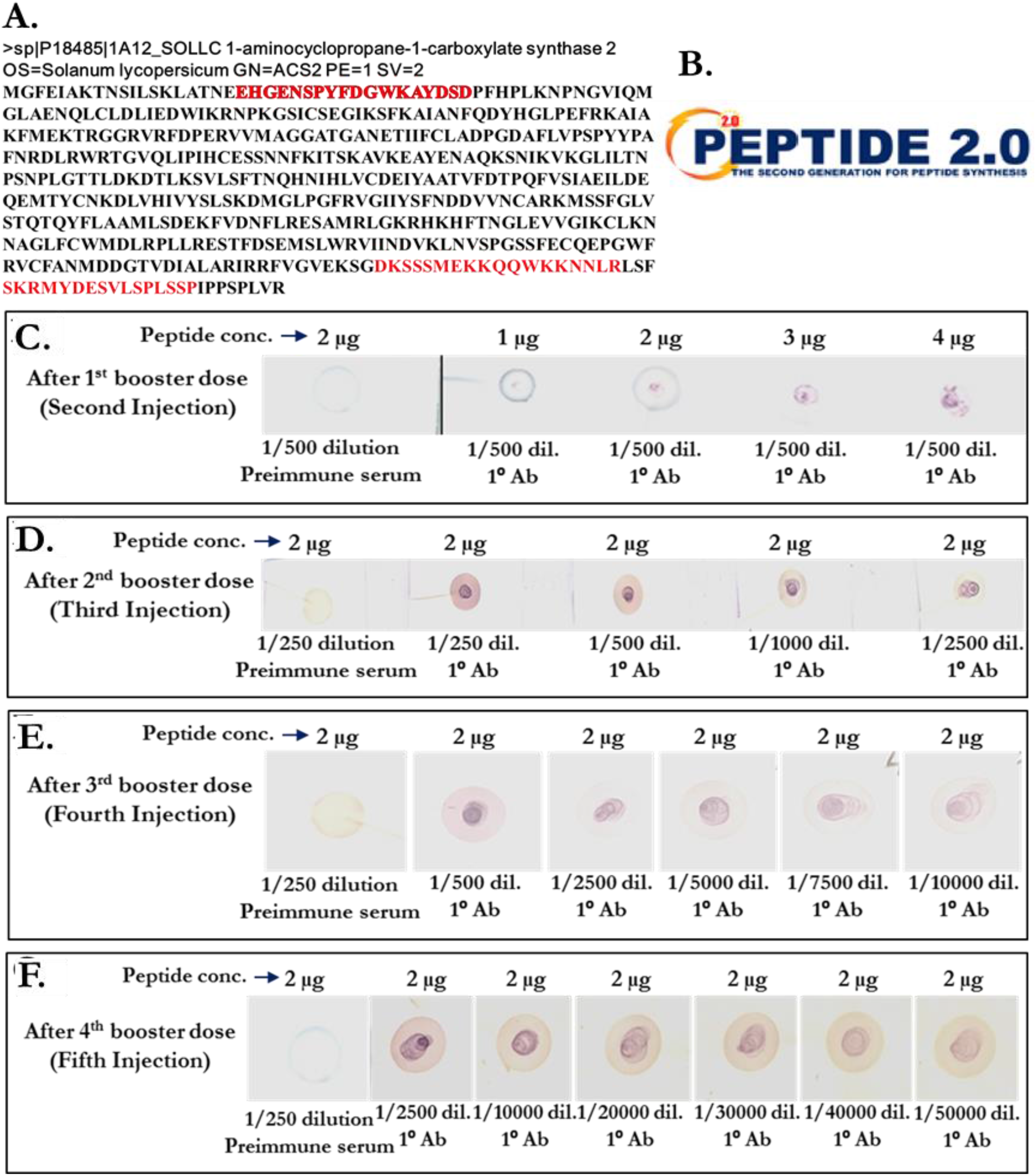
Raising and validation of antibodies specific for ACS2 peptides. Three antigenic peptides (marked in red) specific to ACS2 protein were predicted by PEPTIDE 2.0 (https://www.peptide2.com). Out of three peptides, N-terminal region peptide “EHGENSPYFDGWKAYDSD” specific to ACS2 protein was selected for antibody raising and was synthesized by PEPTIDE 2.0 **(A)**. The small aliquots of antisera from rabbit were collected after 1^st^, 2^nd^, 3^rd^, and 4^th^ booster dose and checked for antibody titer value by DOT BLOT assay **(B-E)**.

**Supplemental Figure S6.**
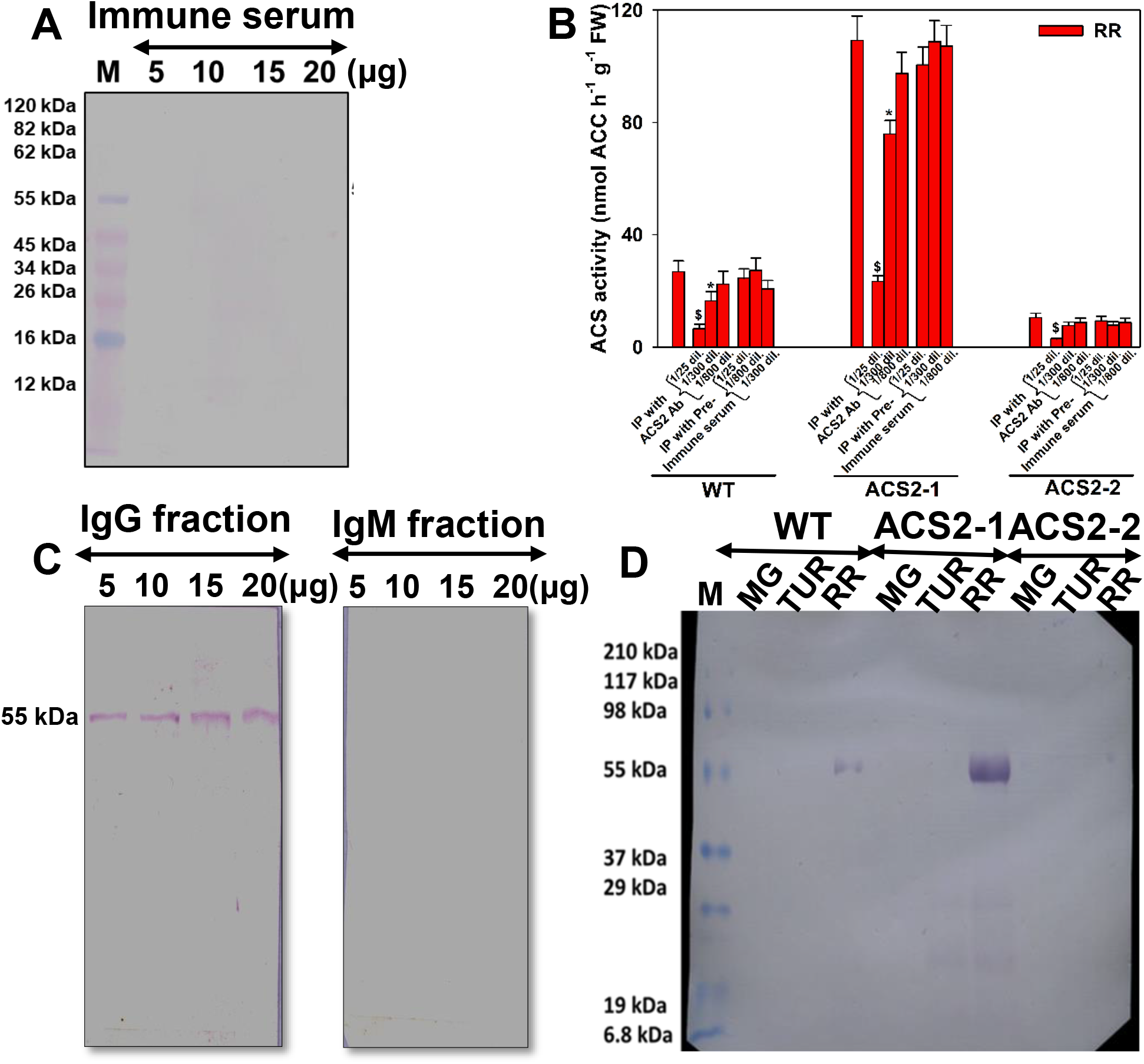
Immunoprecipitation and Western blotting of ACS2 protein using purified IgG fraction. **(A)** Varying amount (5–20 μg) of crude extract of wild-type red-ripe fruits was resolved on SDS-PAGE gel with protein marker (**M**), and Western blotting was performed using unpurified immune serum. **(B)** Immunoprecipitation of ACS protein was carried out in crude extract of wild-type red-ripe fruits with protein-A Sepharose beads. After precipitating antigen-antibody complex, the supernatants were analyzed for the ACS activity. (Student’s t-test; * for P ≤ 0.05, # for P ≤ 0.01 and $ for P ≤ 0.001, for each fruit maturity stage, n=3 ± SE). **(C)** The immunoprecipitated ACS2 protein was resolved on the SDS-PAGE gel. The Western blotting was performed using purified IgG (Left) and IgM fraction (Right) of antiserum. **(D)** Western blot assay of immunoprecipitated ACS2 protein at different stages of tomato fruit ripening. IgG-mediated, immunoprecipitated ACS2 protein content was quantified by Bradford’s assay. An equal amount (20 μg) of immunoprecipitated protein from different ripening stages of tomato fruits of WT and mutants was resolved on SDS-PAGE gel.

**Supplemental Figure S7.**
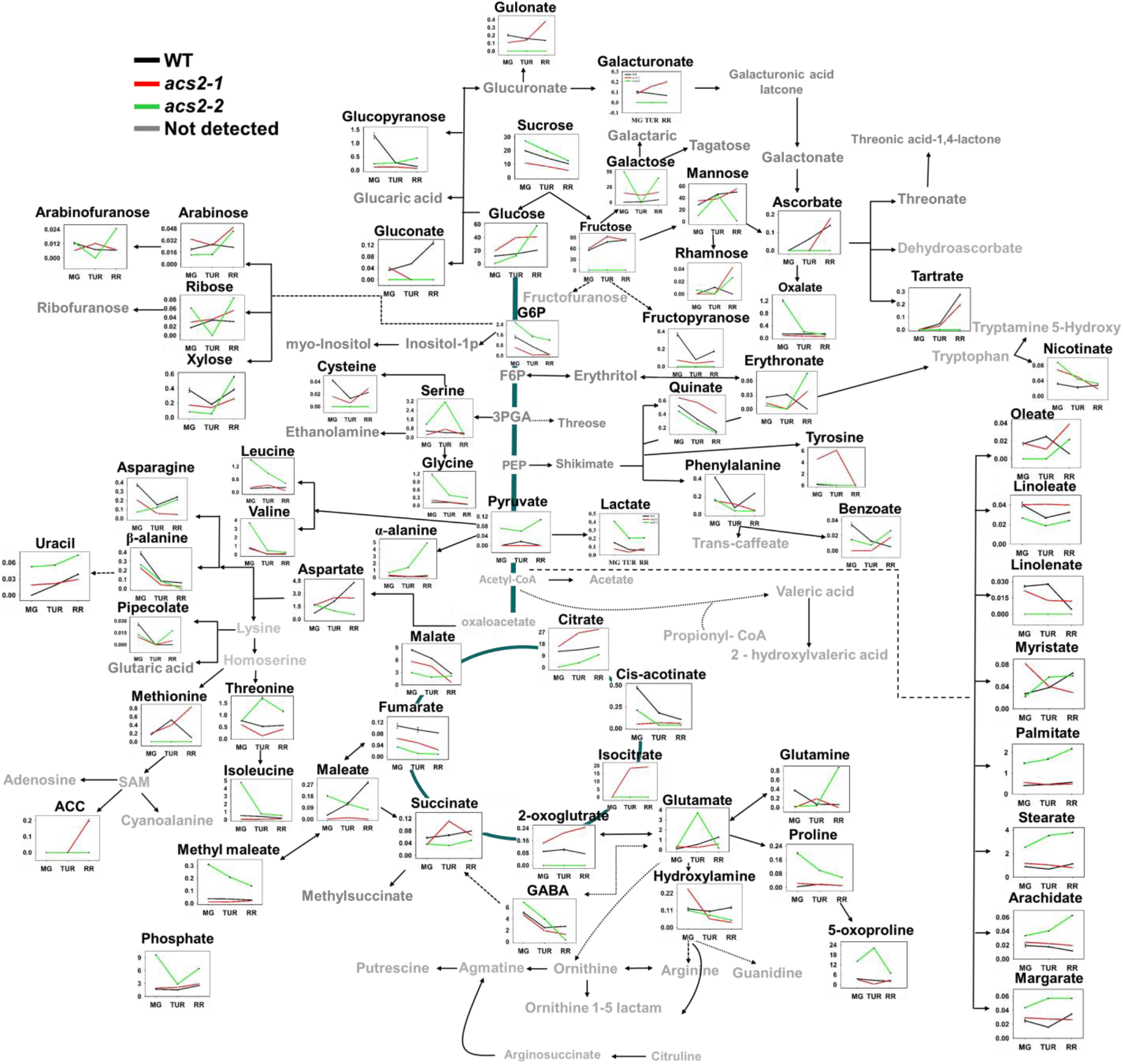
The metabolic shifts in *acs2* mutant fruits during ripening in comparison to WT. An overview of the metabolic pathways representing relative abundance of metabolites at different ripening stages in fruits of WT, *acs2–1*, and *acs2–2* mutants. The Y-axis represents the relative levels of metabolites with reference to internal standard ribitol. X axis denotes metabolite levels at MG-mature green, TUR-turning, and RR-red-ripe stages.

**Supplemental Table S1.** EMS-mutagenized tomato populations used for the isolation of *ACS2* mutants. Four different EMS-mutagenized populations were screened for detection of mutations in *ACS2* gene (Solyc01g095080.2.1) using TILLING. The number of mutagenized plants screened is indicated in the table.

| Population Number | Tomato cultivar | EMS concentration | Generation used for DNA pooling | Population size | Number of *acs-2* mutant alleles identified |
| --- | --- | --- | --- | --- | --- |
| I | Arka Vikas | 60 mM | M<sub>2</sub> | 3768 | 5 |
| II | Arka Vikas | 120 mM | M<sub>2</sub> | 768 | 1 |
| III | M82 | 60 mM | M<sub>2</sub>* | 1536 | 2 |
| IV | M82 | 60 mM | M<sub>3</sub> | 3072 | 1 |

\*The population III was obtained from Dani Zamir, Hebrew University, Israel. The raising of this population is described in Menda et al., (2014). The population IV consisted of M<sub>3</sub> seeds harvested from population III.

**Supplemental Table S2.**
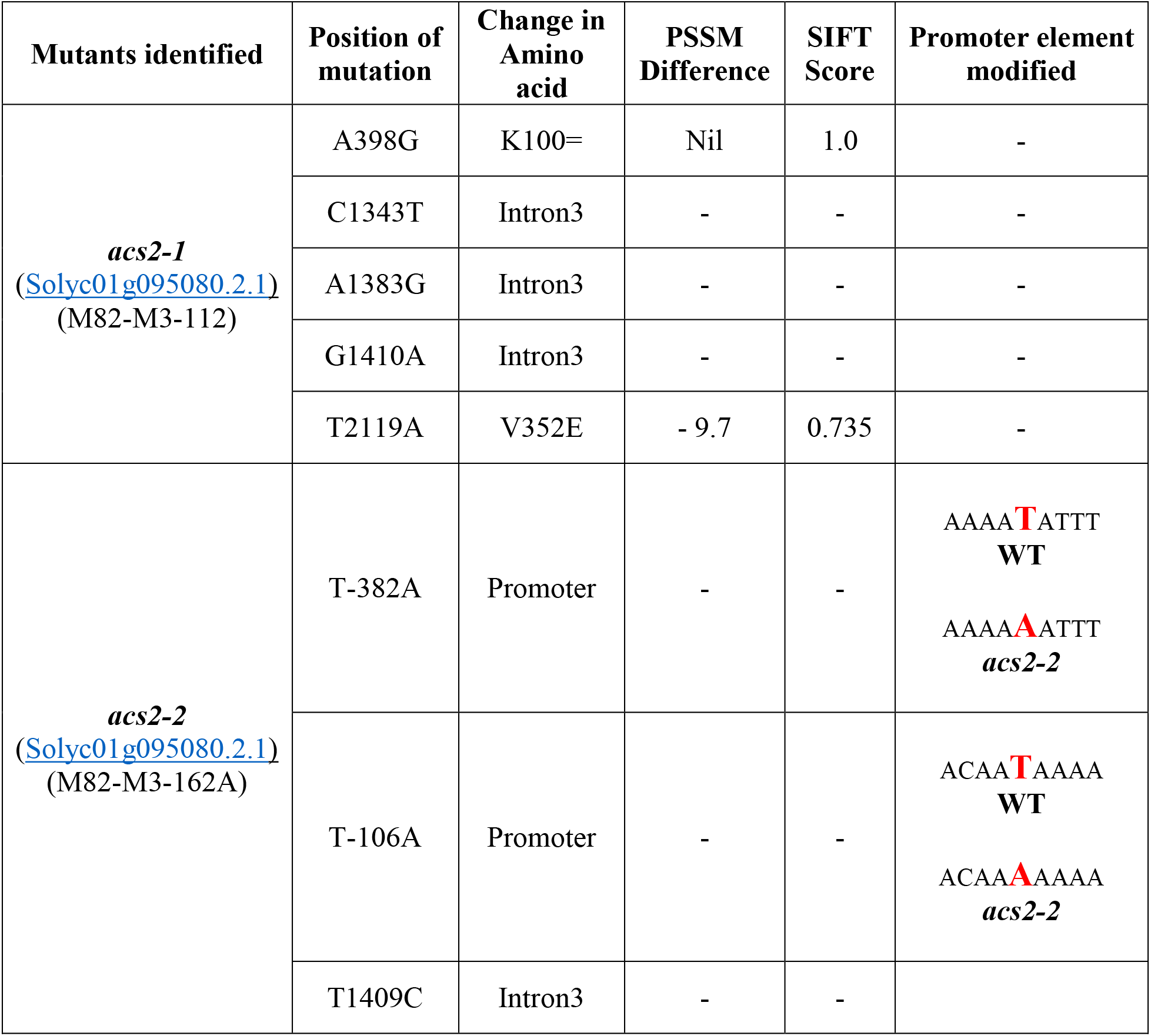
*ACS2* Mutant lines identified and confirmed for mutation. The position of mutation was confirmed by Multialin Interface page (Corpet et al., 1988) using WT DNA as a template sequence. The changes in amino acids were obtained after the translation of mutated ORF sequence, The effect of the non-synonymous mutation on protein function was determined by using SIFT software version 4.0.5 (https://sift.bii.a-star.edu.sg/). The PSSM difference was obtained using NCBI PSSM viewer (https://www.ncbi.nlm.nih.gov/Class/Structure/pssm/pssm_viewer.cgi)

**Supplemental Table S3.** The genetic segregation of *acs2–1* and *acs2–2* mutants in BC1F2 generation. The presence of mutated gene copy and its zygosity in backcrossed plants was monitored by the CEL-I endonuclease assay (Mohan et al., 2016). In brief, we used PCR based mutation screening which involves three steps – firstly, amplification of a fragment of interest followed by heteroduplex formation and mismatch cleavage by CEL-I enzyme and finally, detection on denaturing polyacrylamide gels.

| Crosses | BC<sub>1</sub>F<sub>2</sub><br>Population<br>size | BC<sub>1</sub>F<sub>2</sub><br>(WT<br>homozygous)<br>(+/+) | BC<sub>1</sub>F<sub>2</sub><br>(heterozygous)<br>(+/- ) | BC<sub>1</sub>F<sub>2</sub><br>(mutant<br>homozygous)<br>(- /- ) | χ<sup>2</sup> | P-Value |
| --- | --- | --- | --- | --- | --- | --- |
| *acs2-1*♂<br>X<br>WT♀ | 120 | 30 | 61 | 29 | 0.224<br>(3:1)<br>0.245<br>(1:2:1) | 0.62<br>(3:1)<br>0.88<br>(1:2:1) |
| *acs2-2*♂<br>X<br>WT♀ | 108 | 27 | 56 | 25 | 0.101<br>(3:1)<br>0.113<br>(1:2:1) | 0.95<br>(3:1)<br>0.94<br>(1:2:1) |

**Supplemental Table S4.** Increase in ACS2–1 protein stability predicted by different software. The impact of the mutations in *acs2–1* on the stability of ACS2–1 protein was analyzed using five different software (CUPSAT, MAESTROWeb, PoPMuSiCv3.1, STRUM, and DynaMut). All software predicted that the mutated protein to be more stable than the wild type.

| Tool | Chain A<br>$\Delta\Delta G$<br>(kcal/mol) | Chain B<br>$\Delta\Delta G$<br>(kcal/mol) | Stabilizing/<br>Unstabilizing | Link |
| --- | --- | --- | --- | --- |
| CUPSAT | 0.77 | 0.91 | Stabilizing | http://cupsat.tu-bs.de/ |
| MAESTROWeb | - 0.406 | - 0.406 | Stabilizing | https://pbwww.che.sbg.ac.at/maestro/web |
| PoPMuSiCv3.1 | - 0.34 | - 0.34 | Stabilizing | https://soft.dezyme.com/query/create/pop |
| STRUM | - 0.34 | - 0.34 | Stabilizing | https://zhanglab.ccmb.med.umich.edu/STRUM/ |
| DynaMut | 0.38 | 0.223 | Stabilizing | http://biosig.unimelb.edu.au/dynamut/ |

$\Delta\Delta G$ : The change in Gibbs free energy. The above software predicted that observed changes $\Delta\Delta G$ due to point mutation stabilizes the mutated protein.

**Supplemental Table S5.**
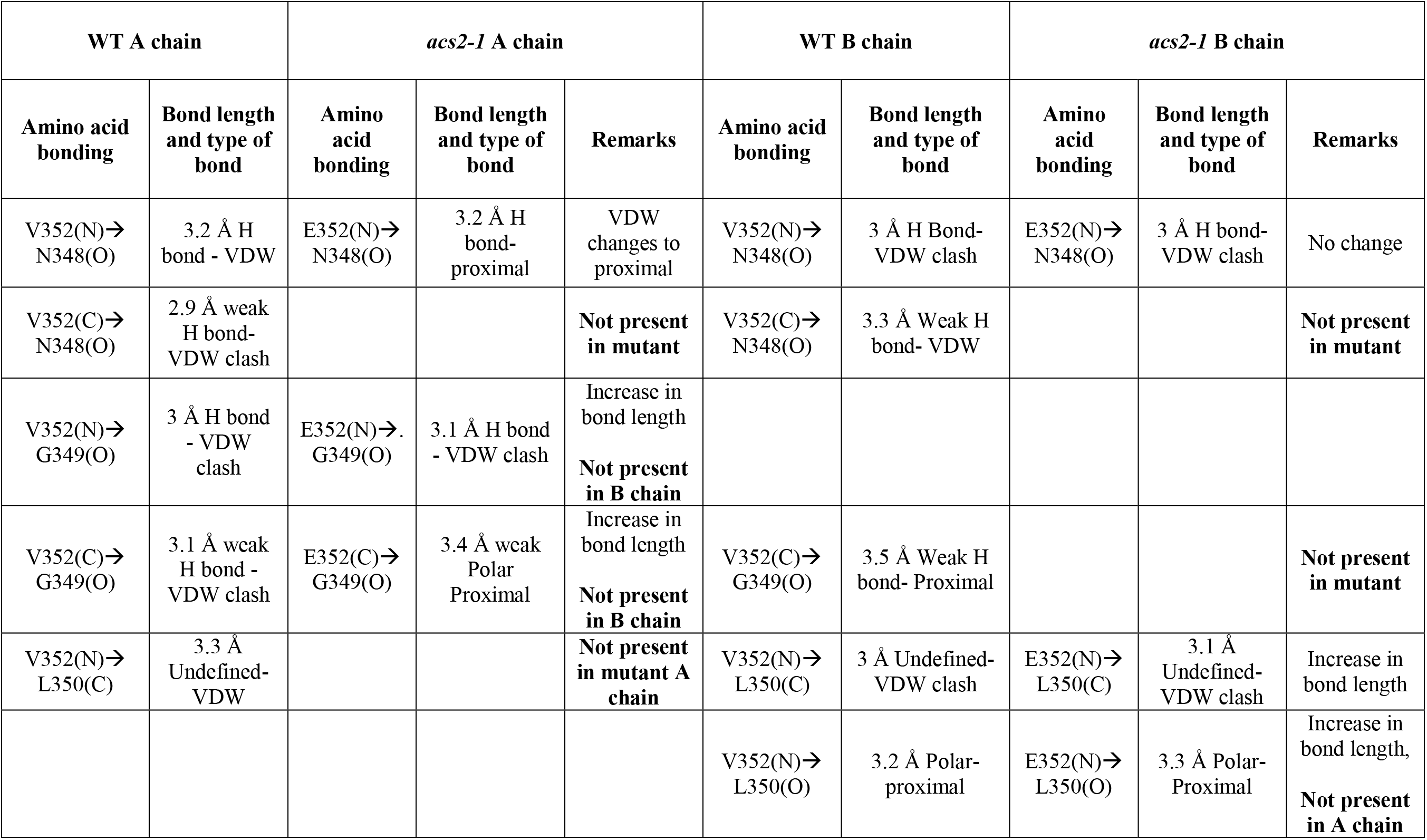
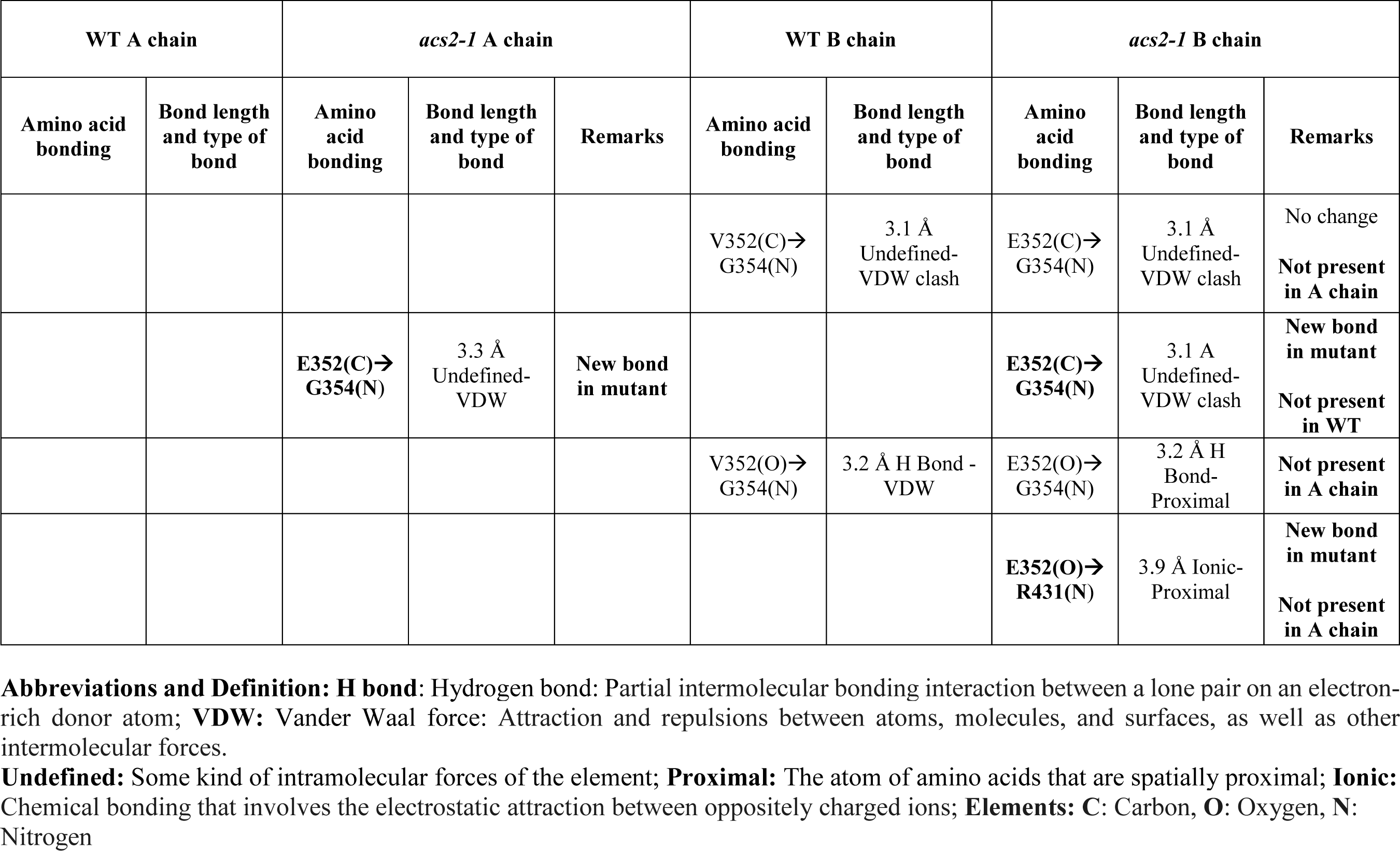
The alteration of bonding pattern in ACS2 protein in *acs2–1* mutant compared to WT protein. The interacting amino acids are denoted by respective alphabets (capital). The bonding atoms in each amino acid are represented in parentheses, and bond length is denoted in angstroms (Å). The novel amino acid interactions appearing in mutant proteins are highlighted in Figure 1B of the main manuscript. The values presented in the table were obtained using Dynamut software (http://biosig.unimelb.edu.au/dynamut/)

| WT A chain |  | *acs2-1* A chain |  |  | WT B chain |  | *acs2-1* B chain |  |  |
| --- | --- | --- | --- | --- | --- | --- | --- | --- | --- |
| Amino acid bonding | Bond length and type of bond | Amino acid bonding | Bond length and type of bond | Remarks | Amino acid bonding | Bond length and type of bond | Amino acid bonding | Bond length and type of bond | Remarks |
| V352(N)→N348(O) | 3.2 Å H bond - VDW | E352(N)→N348(O) | 3.2 Å H bond-proximal | VDW changes to proximal | V352(N)→N348(O) | 3 Å H Bond-VDW clash | E352(N)→N348(O) | 3 Å H bond-VDW clash | No change |
| V352(C)→N348(O) | 2.9 Å weak H bond-VDW clash |  |  | **Not present in mutant** | V352(C)→N348(O) | 3.3 Å Weak H bond- VDW |  |  | **Not present in mutant** |
| V352(N)→G349(O) | 3 Å H bond - VDW clash | E352(N)→G349(O) | 3.1 Å H bond - VDW clash | Increase in bond length<br>**Not present in B chain** |  |  |  |  |  |
| V352(C)→G349(O) | 3.1 Å weak H bond - VDW clash | E352(C)→G349(O) | 3.4 Å weak Polar Proximal | Increase in bond length<br>**Not present in B chain** | V352(C)→G349(O) | 3.5 Å Weak H bond- Proximal |  |  | **Not present in mutant** |
| V352(N)→L350(C) | 3.3 Å Undefined-VDW |  |  | **Not present in mutant A chain** | V352(N)→L350(C) | 3 Å Undefined-VDW clash | E352(N)→L350(C) | 3.1 Å Undefined-VDW clash | Increase in bond length |
|  |  |  |  |  | V352(N)→L350(O) | 3.2 Å Polar-proximal | E352(N)→L350(O) | 3.3 Å Polar-Proximal | Increase in bond length,<br>**Not present in A chain** |
| Amino acid bonding | Bond length and type of bond | Amino acid bonding | Bond length and type of bond | Remarks | Amino acid bonding | Bond length and type of bond | Amino acid bonding | Bond length and type of bond | Remarks |
|  |  |  |  |  | V352(C)→G354(N) | 3.1 Å<br>Undefined-VDW clash | E352(C)→G354(N) | 3.1 Å<br>Undefined-VDW clash | No change<br>**Not present in A chain** |
|  |  | E352(C)→G354(N) | 3.3 Å<br>Undefined-VDW | **New bond in mutant** |  |  | E352(C)→G354(N) | 3.1 Å<br>Undefined-VDW clash | **New bond in mutant**<br>**Not present in WT** |
|  |  |  |  |  | V352(O)→G354(N) | 3.2 Å H Bond - VDW | E352(O)→G354(N) | 3.2 Å H Bond-Proximal | **Not present in A chain** |
|  |  |  |  |  |  |  | E352(O)→R431(N) | 3.9 Å Ionic-Proximal | **New bond in mutant**<br>**Not present in A chain** |

**Abbreviations and Definition:** **H bond:** Hydrogen bond: Partial intermolecular bonding interaction between a lone pair on an electron-rich donor atom; **VDW:** Vander Waal force: Attraction and repulsions between atoms, molecules, and surfaces, as well as other intermolecular forces.
**Undefined:** Some kind of intramolecular forces of the element; **Proximal:** The atom of amino acids that are spatially proximal; **Ionic:** Chemical bonding that involves the electrostatic attraction between oppositely charged ions; **Elements:** C: Carbon, O: Oxygen, N: Nitrogen

**Supplemental Table S6.** Disruption in transcription factor binding site due to promoter mutations. The promoter sequence of the *ACS2* gene was analyzed by Plant ChIP-seq Database (PCBase; http://pcbase.itps.ncku.edu.tw) to predict the sites for binding of transcription factors and other DNA binding proteins. The promoter mutations altered the binding sites of transcription factors and DNA binding proteins listed below. For details, see Supplemental dataset 1.

| Mutated base position on *ACS2* promoter | Binding site in WT | Binding sites in *acs2- 2* mutant | Remarks |
| --- | --- | --- | --- |
| -106 ± 4 | 1. SOC1<br>2. ABF4 | 1. AZF<br>2. ABF4 | Lost SOC1 binding and gained AZF transcription factor binding<br>AZF has strong homology with tomato Zinc finger, C2H2- type transcription factor (Solyc04g077980.1), a putative transcriptional repressor |
| -382 ± 4 | 1. 3-SEP<br>2. ZmCCA1B | 1. 3-SEP<br>2. ZmCCA1B | No change |

**Supplemental Table S7.** Changes in methylation status of *ACS2* promoter at −106 and −382 position during fruit ripening. The methylation data were downloaded from Tomato Epigenome Database (http://ted.bti.cornell.edu/epigenome/index.html). The table lists the methylation status at −106 and −382 position (mutated in *acs2–2*) of ACS2 promotor at different stages of fruit ripening. For the full table see Supplemental dataset 2.

| Mutation<br>Position<br>(ITAG2.5) | Chromosome<br>Position<br>(ITAG2.4) |  | Context | MG |  |  | BR |  |  | BR+10 |  |  | CNR_BR |  |  |
| --- | --- | --- | --- | --- | --- | --- | --- | --- | --- | --- | --- | --- | --- | --- | --- |
|  | Position | Strand |  | 5mc | C | Ratio | 5mc | C | Ratio | 5mc | C | Ratio | 5mc | C | Ratio |
| -106 | 78217288 | + | CHH | 2 | 12 | 14% | 1 | 8 | 11% | 0 | 2 | 0% | 0 | 0 | n/a |
| -382 | 78217565 | - | CWG | 4 | 1 | 80% | 9 | 4 | 69% | 1 | 6 | 14% | 2 | 1 | 66% |

**Supplemental Table S8.** Comparisons of flower numbers and fruit set in WT and *ACS2* mutants. The data was recorded for five independent plants (65-day-old) of each mutant. All flower truss present in 65-day-old plants were considered here.

| **Plant accession** | **No. of flowers/truss** | **Total no. of flowers/**<br>**plant** | **Total no. of fruit**<br>**set/truss** |
| --- | --- | --- | --- |
| WT (M82) | 6-8 | 90-100 | 6-7 |
| *acs2-1* | 7-10 | 60-80 | 6-9 |
| *acs2-1/acs2-1* | 5-7 | 70-90 | 4-6 |
| *acs2-2* | 5-7 | 45-65 | 3-5 |
| *acs2-2/acs2-2* | 5-7 | 45-65 | 3-5 |

**Supplemental Table S9.**
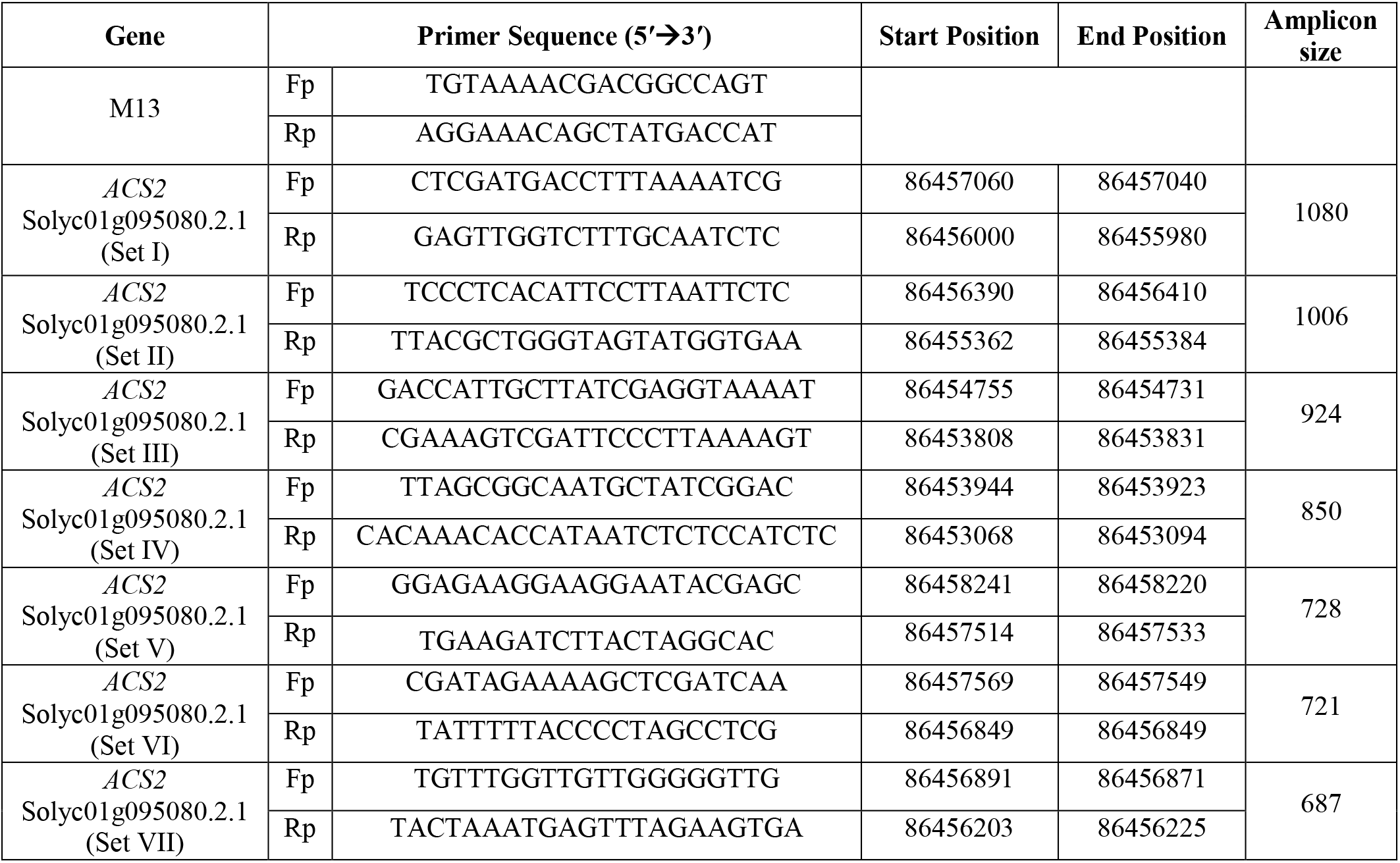
List of primers used for screening for mutations detection in the *ACS2* gene by TILLING. The primers used for screening for mutations by TILLING and Sanger-based sequencing were designed based on the tomato genome sequence SGN SL2.50 version. The *ACS2* gene information and location of the primers on the genome sequence is also indicated. M13 tailed primers were used for screening for the mutations in the *ACS2* gene on Li-COR 4300 DNA analyzer using TILLING.

**Supplemental Table S10.**
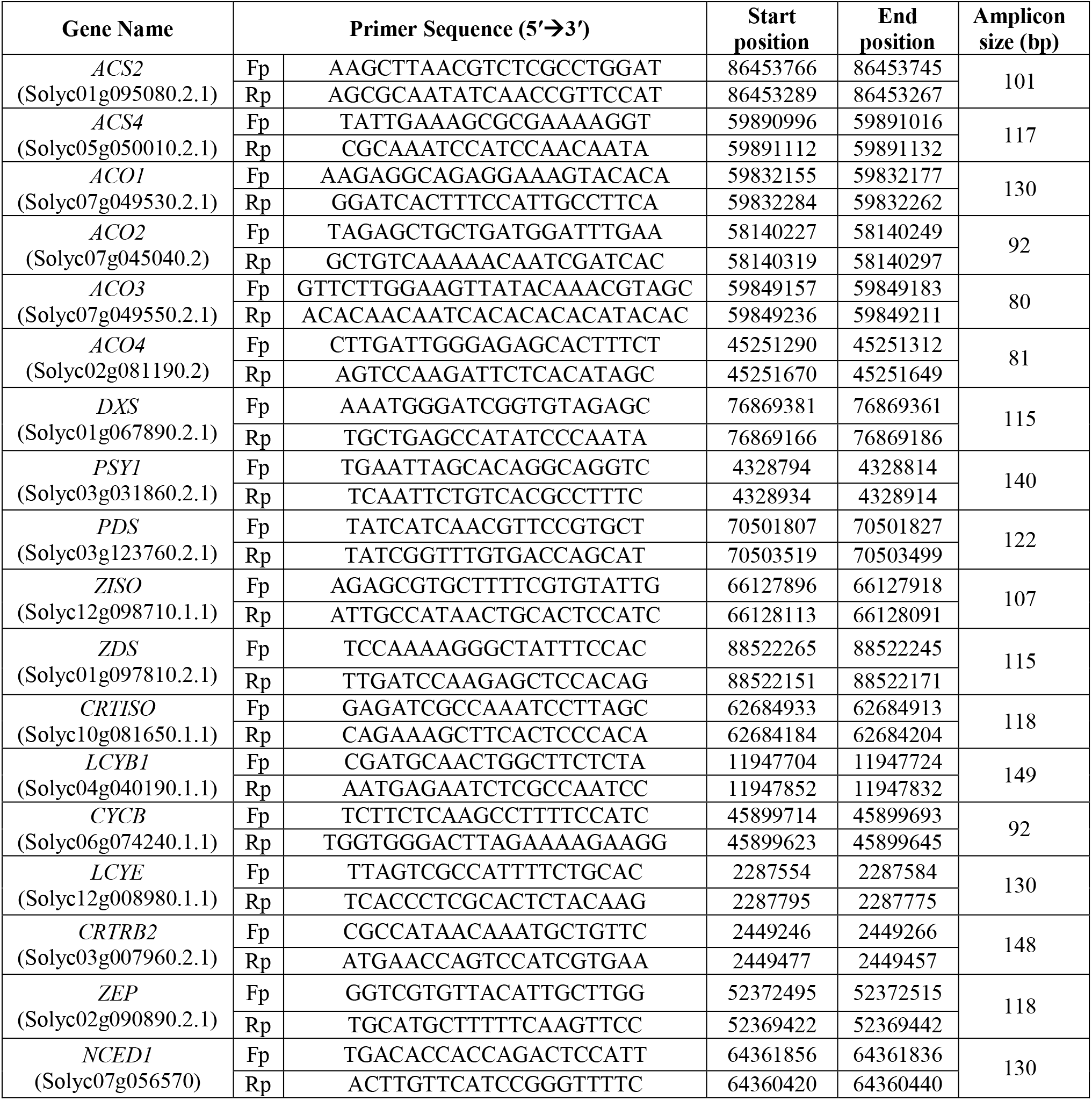
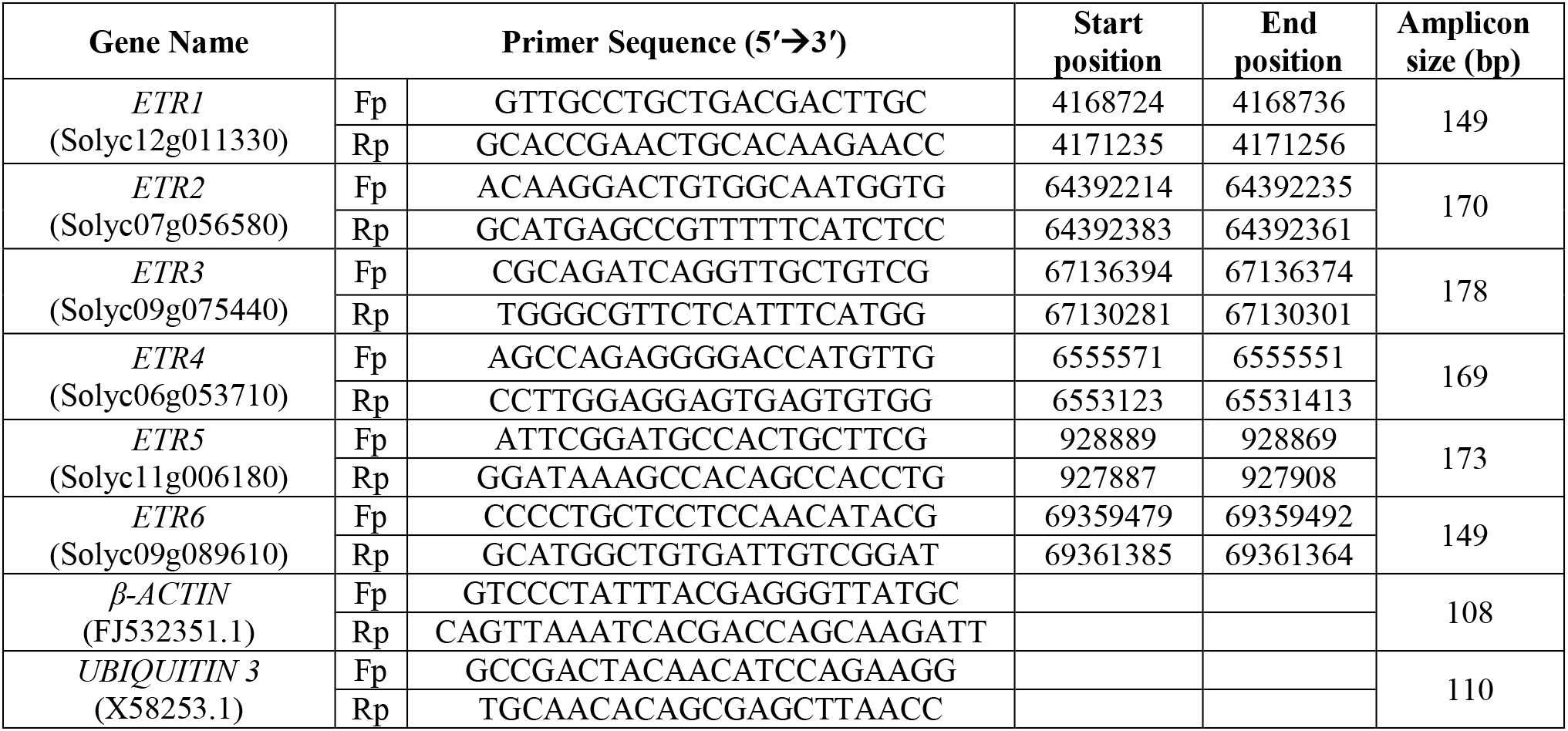
List of genes and the primers used for qRT-PCR analysis. The gene coordinates are according to SGN SL2.50 version. The genes are as follows: *ACS*-*1-aminocyclopropane-1-carboxylate synthase (isoforms 2 and 4); ACO-1-aminocyclopropane-1-carboxylate oxidase (isoforms 1 to 4); DXS-Deoxy-xylulose 5-phosphate synthase, PSY1-Phytoene synthase1, PDS-Phytoene desaturase, ZISO-ζ-carotene isomerase, ZDS-ζ-carotene desaturase; CRTISO-Carotenoid isomerase, LCYB1-Lycopene β-cyclase 1, CYCB-Chromoplast specific Lycopene β-cyclase, LCYE-Lycopene ε-cyclase, CRTRB2-β-carotene hydroxylase 2, ZEP-Zeaxanthin epoxidase, NCED1–9-cis-epoxycarotenoid dioxygenase1, ETR-Ethylene receptor like protein (isoform 1 to 6); β-ACTIN, UBIQUITIN 3.*

## REFERENCES

Abeles FB, Morgan PW, Saltveit Jr ME. (1992) Ethylene in plant biology. Academic press

Adams DO, Yang SF. (1979) Ethylene biosynthesis: identification of 1-aminocyclopropane-1-carboxylic acid as an intermediate in the conversion of methionine to ethylene. Proc. Natl. Acad. Sci. USA. 76: 170–174.

Alba R, Payton P, Fei Z, McQuinn R, Debbie P, Martin GB, Tanksley SD, Giovannoni JJ. (2005) Transcriptome and selected metabolite analyses reveal multiple points of ethylene control during tomato fruit development. Plant Cell. 17: 2954–2965.

Barry CS. (2014) Ripening mutants. *In* “Fruit Ripening: Physiology, Signalling and Genomics” Eds. Nath P, Bouzayen M, Mattoo AK, Pech JC, CABI; pp 246–258

Barry CS, Blume B, Bouzayen M, Cooper W, Hamilton AJ, Grierson D. (1996) Differential expression of the 1-aminocyclopropane-1-carboxylate oxidase gene family of tomato. Plant J. 9: 525–535.

Barry CS, Fox EA, Yen HC, Lee S, Ying TJ, Grierson D, Giovannoni JJ. (2001) Analysis of the Ethylene Response in the epinastic Mutant of Tomato. Plant Physiol.127: 58–66.

Barry CS, Llop-Tous MI, Grierson D. (2000) The regulation of 1-aminocyclopropane-1-carboxylic acid synthase gene expression during the transition from system-1 to system-2 ethylene synthesis in tomato. Plant Physiol. 123: 979–986.

Bastías A, Yañez M, Osorio S, Arbona V, Gómez-Cadenas A, Fernie AR, Casaretto JA. (2014) The transcription factor AREB1 regulates primary metabolic pathways in tomato fruits. J. Exp. Bot. 65: 2351–2363.

Bleecker AB, Kenyon WH, Somerville SC, Kende H. (1986) Use of monoclonal antibodies in the purification and characterization of 1-aminocyclopropane-1-carboxylic acid synthase, an enzyme in ethylene biosynthesis. Proc. Natl. Acad. Sci. USA 83:7755–7759.

Bodanapu R, Gupta SK, Basha PO, Sakthivel K, Sadhna, Sreelakshmi Y, Sharma R. (2016) Nitric oxide overproduction in tomato *shr* mutant shifts metabolic profiles and suppresses fruit growth and ripening. Front. Plant Sci. 7: 1714

Bradford MM. (1976) A rapid and sensitive method for the quantitation of microgram quantities of protein utilizing the principle of protein-dye binding. Anal. Biochem. 72: 248–254.

Brady CJ. (1987) Fruit ripening. Annu. Rev. Plant Physiol. 38: 155–178

Breitel DA, Chappell-Maor L, Meir S, Panizel I, Puig CP, Hao Y, Yifhar T, Yasuor H, Zouine M, Bouzayen M, Richart AG, Rogachev I, Aharoni A. (2016) AUXIN RESPONSE FACTOR 2 intersects hormonal signals in the regulation of tomato fruit ripening. PLoS Genet. 12(3): e1005903.

Bulens I, Van de Poel B, Hertog ML, De Proft MP, Geeraerd AH, Nicolaï BM. (2011) Protocol: an updated integrated methodology for analysis of metabolites and enzyme activities of ethylene biosynthesis. Plant Methods. 7: 17.

Chae HS, Faure F, Kieber JJ. (2003) The eto1, eto2, and eto3 mutations and cytokinin treatment increase ethylene biosynthesis in Arabidopsis by increasing the stability of ACS protein. Plant Cell. 15: 545–559.

Chae HS, Kieber JJ. (2005) Eto Brute? Role of ACS turnover in regulating ethylene biosynthesis. Trends Plant Sci. 10: 291–296.

Chen G, Alexander L, Grierson D. (2004). Constitutive expression of EIL-like transcription factor partially restores ripening in the ethylene-insensitive Nr tomato mutant. J. Exp. Bot. 55:1491–1497.

Cooper AJ. (2004) The role of glutamine transaminase K (GTK) in sulfur and alpha-keto acid metabolism in the brain, and in the possible bioactivation of neurotoxicants. Neurochem. Int. 44, 557–577.

Duan J, Shi J, Ge X, Dölken L, Moy W, He D, Shi S, Sanders AR, Ross J, Gejman PV. (2013) Genome-wide survey of inter-individual differences of RNA stability in human lymphoblastoid cell lines. Sci. Rep. 20:1318

Ellens KW, Richardson LG, Frelin O, Collins J, Ribeiro CL, Hsieh YF, Mullen RT, Hanson AD. (2015) Evidence that glutamine transaminase and omega-amidase potentially act in tandem to close the methionine salvage cycle in bacteria and plants. Phytochem. 113: 160–169.

Fernie AR, Tohge T. (2017) The genetics of plant metabolism. Annu. Rev. Genet. 51: 287–310.

Fujino DW, Burger DW, Bradford KJ. (1989) Ineffectiveness of ethylene biosynthetic and action inhibitors in phenotypically reverting the Epinastic mutant of Tomato (Lycopersicon esculentum mill.). J. Plant Growth Regul. 8: 53–61.

Fujisawa M, Nakano T, Shima Y, Ito Y. (2013) A large-scale identification of direct targets of the tomato MADS box transcription factor RIPENING INHIBITOR reveals the regulation of fruit ripening. Plant Cell. 25: 371–386.

Fujisawa M, Shima Y, Higuchi N, Nakano T, Koyama Y, Kasumi T, Ito Y. (2012) Direct targets of the tomato-ripening regulator RIN identified by transcriptome and chromatin immunoprecipitation analyses. Planta. 235: 1107–1122.

Ghanem ME, Albacete A, Martínez-Andújar C, Acosta M, Romero-Aranda R, Dodd IC, Lutts S, Pérez-Alfocea F. (2008) Hormonal changes during salinity-induced leaf senescence in tomato (Solanum lycopersicum L.). J. Exp. Bot. 59: 3039–3050.

Gupta KJ, Shah JK, Brotman Y, Jahnke K, Willmitzer L, Kaiser WM, Bauwe H, Igamberdiev AU. (2012) Inhibition of aconitase by nitric oxide leads to induction of the alternative oxidase and to a shift of metabolism towards biosynthesis of amino acids. J. Exp. Bot. 63: 1773–1784.

Gupta P, Sreelakshmi Y, Sharma R. (2015) A rapid and sensitive method for determination of carotenoids in plant tissues by high performance liquid chromatography. Plant Methods. 11: 5

Gupta SK, Sharma S, Santisree P, Kilambi HV, Appenroth K, Sreelakshmi Y, Sharma R. (2014) Complex and shifting interactions of phytochromes regulate fruit development in tomato. Plant Cell Environ. 37:1688–1702

Guzman P, Ecker JR. (1990) Exploiting the triple response of Arabidopsis to identify ethylene-related mutants. Plant Cell. 2: 513–523.

Hackett RM, Ho C-W, Lin Z, Foote HCC, Fray RG, Grierson D. (2000) Antisense inhibition of the Nr gene restores normal ripening to the tomato Never-ripe mutant, consistent with the ethylene receptor-inhibition model. Plant Physiol. 124: 1079–1086.

Hamilton AJ, Lycett GW, Grierson D. (1990). Antisense gene that inhibits synthesis of the hormone ethylene in transgenic plants. Nature 346: 284–287.

Hebsgaard SM, Korning PG, Tolstrup N, Engelbrecht J, Rouze P, Brunak S. (1996) Splice site prediction in Arabidopsis thaliana DNA by combining local and global sequence information, Nucleic Acids Res. 24: 3439–3452

Hoffman NE, Yang SF, McKeon T. (1982) Identification of 1-(malonylamino) cyclopropane-1-carboxylic acid as a major conjugate of 1-aminocyclopropane-1-carboxylic acid, an ethylene precursor in higher plants. Biochem. Bioph. Res. Co. 104: 765–770.

Hoffman NE, Yang SF. (1980) Changes of 1-aminocyclopropane-1-carboxylic acid content in ripening fruits in relation to their ethylene production rates. J. Am. Soc. Hortic. Sci. 105: 492–495.

Huai Q, Xia Y, Chen Y, Callahan B, Li N, Ke H. (2001) Crystal structures of 1-aminocyclopropane-1-carboxylate (ACC) synthase in complex with amino ethoxy-vinylglycine and pyridoxal-5′-phosphate provide new insight into catalytic mechanisms. J. Biol. Chem. 276:38210–38216

Itkin M, Seybold H, Breitel D, Rogachev I, Meir S, Aharoni A. (2009) TOMATO AGAMOUS-LIKE 1 is a component of the fruit ripening regulatory network. Plant J. 60: 1081–1095.

Ito Y, Kitagawa M, Ihashi N, Yabe K, Kimbara J, Yasuda J, Ito H, Inakuma T, Hiroi S, Kasumi T. (2008) DNA-binding specificity, transcriptional activation potential, and the rin mutation effect for the tomato fruit-ripening regulator RIN. Plant J. 55: 212–223.

Ito Y, Nishizawa-Yokoi A, Endo M, Mikami M, Shima Y, Nakamura N, Kotake-Nara E, Kawasaki S, Toki S. (2017) Re-evaluation of the rin mutation and the role of RIN in the induction of tomato ripening. Nature Plants. 3:866–874.

Jia C, Zhang L, Liu L, Wang J, Li C, Wang Q. (2013) Multiple phytohormone signalling pathways modulate susceptibility of tomato plants to *Alternaria alternata* f. sp. *lycopersici*. J. Exp. Bot. 64: 637–650.

John I, Drake R, Farrell A, Cooper W, Lee P, Horton P, Grierson D. (1995) Delayed leaf senescence in ethylene-deficient ACC-oxidase antisense tomato plants: molecular and physiological analysis. Plant J. 7: 483–490.

Kamiyoshihara Y, Iwata M, Fukaya T, Tatsuki M, Mori H. (2010) Turnover of *LeACS2*, a wound inducible 1-aminocyclopropane-1-carboxylic acid synthase in tomato, is regulated by phosphorylation/dephosphorylation. Plant J. 64, 140–150.

Kevany BM, Tieman DM, Taylor MG, Cin VD, Klee HJ. (2007) Ethylene receptor degradation controls the timing of ripening in tomato fruit. Plant J. 51: 458–467.

Kieber JJ, Rothenberg M, Roman G, Feldmann KA, Ecker JR. (1993). CTR1, a negative regulator of the ethylene response pathway in Arabidopsis, encodes a member of the raf family of protein kinases. Cell 72: 427–441.

Kilambi HV, Kumar R, Sharma R, Sreelakshmi Y. (2013) Chromoplast-specific carotenoid-associated protein appears to be important for enhanced accumulation of carotenoids in *hp1* tomato fruits. Plant Physiol. 161:2085–2101

Klee HJ, Giovannoni JJ. (2011) Genetics and control of tomato fruit ripening and quality attributes. Annu. Rev. Genet. 45: 41–59.

Kodaira KS, Qin F, Tran LS, Maruyama K, Kidokoro S, Fujita Y, Shinozaki K, Yamaguchi-Shinozaki K. (2011) Arabidopsis Cys2/His2 zinc-finger proteins AZF1 and AZF2 negatively regulate abscisic acid-repressive and auxin-inducible genes under abiotic stress conditions. Plant Physiol. 157: 742–756

Laemmli UK. (1970) Cleavage of structural proteins during the assembly of the head of bacteriophage T4. Nature. 227: 680–685.

Lanahan MB, Yen HC, Giovannoni JJ, Klee HJ. (1994) The never ripe mutation blocks ethylene perception in tomato. Plant Cell 6: 521–530.

Lashbrook CC, Tieman DM, Klee HJ. (1998) Differential regulation of the tomato ETR gene family throughout plant development. Plant J. 15: 243–252.

Lee KY, Baden C, Howie WJ, Bedbrook J, Dunsmuir P. (1997) Post-transcriptional gene silencing of ACC synthase in tomato results from cytoplasmic RNA degradation. Plant J. 12: 1127–1137.

Lincoln JE, Fischer RL. (1988) Regulation of gene expression by ethylene in wild-type and rin tomato (Lycopersicon esculentum) fruit. Plant Physiol. 88: 370–374

Liu L, Wei J, Zhang M, Zhang L, Li C, Wang Q. (2012) Ethylene independent induction of lycopene biosynthesis in tomato fruits by jasmonates. J. Exp. Bot. 63: 5751–5761

Liu M, Pirrello J, Chervin C, Roustan JP, Bouzayen M. (2015) Ethylene control of fruit ripening: revisiting the complex network of transcriptional regulation. Plant Physiol. 169: 2380–2390

Martínez-Romero D, Bailén G, Serrano M, Guillén F, Valverde JM, Zapata P, Castillo S, Valero D. (2007) Tools to maintain postharvest fruit and vegetable quality through the inhibition of ethylene action: a review. Crit. Rev. Food Sci. Nutr. 47: 543–560.

Menda N, Semel Y, Peled D, Eshed Y, Zamir D. (2004) In silico screening of a saturated mutation library of tomato. Plant J. 38: 861–872.

Mohan V, Gupta S, Thomas S, Mickey H, Charakana C, Chauhan VS, Sharma K, Kumar R, Tyagi K, Sarma S, Gupta SK, Kilambi HV, Nongmaithem S, Kumari A, Gupta P, Sreelakshmi Y, Sharma R. (2016) Tomato fruits show wide phenomic diversity but fruit developmental genes show low genomic diversity. PLoS One 11: e0152907.

Mubarok S, Hoshikawa K, Okabe Y, Yano R, Tri MD, Ariizumi T, Ezura H. (2019) Evidence of the functional role of the ethylene receptor genes SlETR4 and SlETR5 in ethylene signal transduction in tomato. Mol. Genet. Genomics. 294: 301–313.

Muday GK, Rahman A, Binder BM. (2012) Auxin and ethylene: collaborators or competitors?. Trends Plant Sci. 17: 181–195.

Nakatsuka A, Murachi S, Okunishi H, Shiomi S, Nakano R, Kubo Y, Inaba A. (1998) Differential expression and internal feedback regulation of 1-aminocyclopropane-1-carboxylate synthase, 1-aminocyclopropane-1-carboxylate oxidase, and ethylene receptor genes in tomato fruit during development and ripening. Plant Physiol. 118: 1295–1305.

Negi S, Sukumar P, Liu X, Cohen JD, Muday GK. (2010) Genetic dissection of the role of ethylene in regulating auxin-dependent lateral and adventitious root formation in tomato. Plant J. 61: 3–15.

Nogueira M, Mora L, Enfissi EM, Bramley PM, Fraser PD. (2013) Subchromoplast sequestration of carotenoids affects regulatory mechanisms in tomato lines expressing different carotenoid gene combinations. Plant Cell 25: 4560–4579.

Oeller PW, Lu MW, Taylor LP, Pike DA, Theologis A. (1991). Reversible inhibition of tomato fruit senescence by antisense RNA. Science. 254: 437–439.

Okabe Y, Asamizu E, Saito T, Matsukura C, Ariizumi T, Brès C, Rothan C, Mizoguchi T, Ezura H. (2011) Tomato TILLING technology: development of a reverse genetics tool for the efficient isolation of mutants from Micro-Tom mutant libraries. Plant Cell Physiol. 52: 1994–2005

Osorio S, Alba R, Damasceno CM, Lopez-Casado G, Lohse M, Zanor MI, Tohge T, Usadel B, Rose JK, Fei Z, Giovannoni JJ, Fernie AR. (2011) Systems biology of tomato fruit development: combined transcript, protein, and metabolite analysis of tomato transcription factor (nor, rin) and ethylene receptor (Nr) mutants reveals novel regulatory interactions. Plant Physiol. 157: 405–425.

Pan X, Welti R, Wang X. (2010) Quantitative analysis of major plant hormones in crude plant extracts by high performance liquid chromatography-mass spectrometry. Nature Protocols. 5: 986–992.

Pirrello J, Jaimes-Miranda F, Sanchez-Ballesta MT, Tournier B, Khalil-Ahmad Q, Regad F, Latche A, Pech JC, Bouzayen M. (2006) Sl-ERF2, a tomato ethylene response factor involved in ethylene response and seed germination Plant Cell Physiol. 47: 1195–1205.

Qin G, Wang Y, Cao B, Wang W, Tian S. (2012) Unraveling the regulatory network of the MADS box transcription factor RIN in fruit ripening. Plant J. 70: 243–255.

Roessner U, Wagner C, Kopka J, Trethewey RN, Willmitzer L. (2000) Simultaneous analysis of metabolites in potato tuber by gas chromatography-mass spectrometry. Plant J. 23: 131–142.

Rottmann WH, Peter GF, Oeller PW, Keller JA, Shen NF, Nagy BP, Taylor LP, Campbell AD, Theologis A. (1991) 1-aminocyclopropane-1-carboxylate synthase in tomato is encoded by a multigene family whose transcription is induced during fruit and floral senescence. J. Mol. Biol. 222:937–961.

Santisree P, Nongmaithem S, Vasuki H, Sreelakshmi Y, Ivanchenko MG, Sharma R. (2011) Tomato Root Penetration in Soil Requires a Co-action between Ethylene and Auxin Signaling. Plant Physiol. 156:1424–1438

Shu K, Liu XD, Xie Q, He ZH. (2016) Two faces of one seed: hormonal regulation of dormancy and germination. Mol. Plant. 9: 34–45.

Sreelakshmi Y, Gupta S, Bodanapu R, Chauhan VS, Hanjabam M, Thomas S, Mohan V, Sharma S, Srinivasan R, Sharma R. (2010) NEATTILL: A simplified procedure for nucleic acid extraction from arrayed tissue for TILLING and other high-throughput reverse genetic applications. Plant Methods 6:3

Su L, Diretto G, Purgatto E, Danoun S, Zouine M, Li Z, Roustan JP, Bouzayen M, Giuliano G, Chervin C. (2015) Carotenoid accumulation during tomato fruit ripening is modulated by the auxin-ethylene balance. BMC Plant Biol. 15: 114.

Theologis A, Oeller PW, Wong LM, Rottmann WH, Gantz DM. (1993) Use of a tomato mutant constructed with reverse genetics to study fruit ripening, a complex developmental process. Dev. Genet. 14: 282–295.

Tieman DM, Taylor MG, Ciardi JA, Klee HJ. (2000) The tomato ethylene receptors NR and LeETR4 are negative regulators of ethylene response and exhibit functional compensation within a multigene family. Proc. Natl. Acad. Sci. USA. 97: 5663–5668.

Tígchelaar EC, Mcglasson WB, Franklin MJ. (1978) Natural and ethephon-stimulated ripening of F1 hybrids of the ripening inhibitor (rin) and non-ripening (nor) mutants of tomato (Lycopersicon esculentum Mill.). Aust. J. Plant Physiol. 5:449–456.

Till BJ, Zerr T, Comai L, Henikoff S. (2006) A protocol for TILLING and Ecotilling in plants and animals. Nature Protocols. 1: 2465–2477.

Towbin H, Staehelin T, Gordon J. (1979) Electrophoretic transfer of proteins from polyacrylamide gels to nitrocellulose sheets: procedure and some applications. Proc. Natl. Acad. Sci. USA.76: 4350–4354.

Van de Poel B, Bulens I, Hertog ML, Nicolai BM, Geeraerd AH. (2014) A transcriptomics-based kinetic model for ethylene biosynthesis in tomato (Solanum lycopersicum) fruit: development, validation and exploration of novel regulatory mechanisms. New Phytol. 202: 952–963.

Van de Poel B, Bulens I, Markoula A, Hertog ML, Dreesen R, Wirtz M, Vandoninck S, Oppermann Y, Keulemans J, Hell R, Waelkens E. (2012) Targeted systems biology profiling of tomato fruit reveals coordination of the Yang cycle and a distinct regulation of ethylene biosynthesis during postclimacteric ripening. Plant Physiol. 160: 1498–1514.

Vrebalov J, Ruezinsky D, Padmanabhan V, White R, Medrano D, Drake R, Schuch W, Giovannoni J. (2002) A MADS-box gene necessary for fruit ripening at the tomato ripening-inhibitor (rin) locus. Science. 296: 343–346.

Wilkinson JQ, Lanahan MB, Yen HC, Giovannoni JJ, Klee HJ. (1995) An ethylene-inducible component of signal transduction encoded by Never-ripe. Science. 270: 1807–1809

Yang SF, Hoffman NE. (1984) Ethylene biosynthesis and its regulation in higher plants. Annu. Rev. Plant Physiol. 35:155–189

Yang Y-X, Ahammed G, Wu C, Fan S, Zhou Y-H. (2015). Crosstalk among Jasmonate, Salicylate and Ethylene Signaling Pathways in Plant Disease and Immune Responses. Curr. Protein Pept. Sc. 16:450–461.

Zhang M, Yuan B, Leng P. (2009) The role of ABA in triggering ethylene biosynthesis and ripening of tomato fruit. J. Exp. Bot. 60: 1579–1588.

Zhong S, Fei Z, Chen YR, Zheng Y, Huang M, Vrebalov J, McQuinn R, Gapper N, Liu B, Xiang J, Shao Y, Giovannoni JJ. (2013) Single-base resolution methylomes of tomato fruit development reveal epigenome modifications associated with ripening. Nature Biotechnol. 31: 154–159.

## Reference

Menda N, Semel Y, Peled D, Eshed Y, Zamir D. (2004) In silico screening of a saturated mutation library of tomato. The Plant Journal. 38: 861–872.

Corpet F, Gouzy J, Kahn D. (1998) The ProDom database of protein domain families. Nucleic Acids Research. 26: 323–326.

Mohan V, Gupta S, Thomas S, Mickey H, Charakana C, Chauhan VS, Sharma K, Kumar R, Tyagi K, Sarma S, Gupta SK, Kilambi HV, Nongmaithem S, Kumari A, Gupta P, Sreelakshmi Y, Sharma R (2016) Tomato fruits show wide phenomic diversity but fruit developmental genes show low genomic diversity. PLoS One 11: e0152907.

Immink RG, Posé D, Ferrario S, Ott F, Kaufmann K, Valentim FL, De Folter S, Van der Wal F, van Dijk AD, Schmid M, Angenent GC. (2012) Characterization of SOC1’s central role in flowering by the identification of its upstream and downstream regulators. Plant Physiology 160: 433–449.

Kodaira KS, Qin F, Tran LS, Maruyama K, Kidokoro S, Fujita Y, Shinozaki K, Yamaguchi-Shinozaki K. (2011) Arabidopsis Cys2/His2 zinc-finger proteins AZF1 and AZF2 negatively regulate abscisic acid-repressive and auxin-inducible genes under abiotic stress conditions. Plant Physiology 157: 742–756.

